# The divergent responses of salinity generalists to hyposaline stress provide insights into the colonization of freshwaters by diatoms

**DOI:** 10.1101/2024.06.02.597024

**Authors:** Kathryn J. Judy, Eveline Pinseel, Kala M. Downey, Jeffrey A. Lewis, Andrew J. Alverson

## Abstract

Environmental transitions, such as the salinity divide separating marine and fresh waters, shape biodiversity over both shallow and deep timescales, opening up new niches and creating opportunities for accelerated speciation and adaptive radiation. Understanding the evolutionary genetic underpinnings behind habitat transitions is therefore a central question in evolutionary biology. We used time-resolved transcriptomics to contrast the hyposalinity stress responses of two ecologically important diatoms: *Skeletonema marinoi* has a deep marine ancestry but recently invaded brackish waters, whereas *Cyclotella cryptica* has deep freshwater ancestry and can withstand a much broader salinity range. *S. marinoi* is less adept at mitigating even mild salinity stress compared to *C. cryptica*, which has distinct mechanisms for rapid mitigation of hyposaline stress and long-term growth in low salinity. We show that the cellular mechanisms underlying low salinity tolerance, which has allowed diversification across freshwater habitats worldwide, includes elements that are both conserved and variable across the diatom lineage. The balance between ancestral and lineage-specific environmental responses in phytoplankton have likely shaped marine–freshwater transitions on evolutionary timescales and, on contemporary timescales, will likely determine which lineages survive and adapt to changing ocean conditions.

## INTRODUCTION

Environmental transitions are often landmark events in evolution^1–3^. On shallow timescales, the colonization of new environments can trigger rapid adaptive evolution, opening up new niches and creating the conditions for ecological speciation^4^. Played out over macroevolutionary timescales, these processes can lead to adaptive radiations^5^ or increases in the rate of speciation^6–8^. Phenotypic plasticity can play a key role in the colonization of new habitats and, once there, directional selection can tailor the phenotype to the new environment, allowing the population to become permanently established^9^. Historical patterns of habitat shifts can be reconstructed through phylogenetics, but a full understanding of how environmental barriers are crossed requires direct observations from genetics or controlled experiments^10,11^. Pushed to their physiological limits, generalists are natural candidates for experimentation to determine how species successfully colonize, become established, and eventually diversify within new habitats.

Diatoms are microalgae found throughout marine and freshwaters, where they play keystone roles in food webs and nutrient cycles. Diatoms are ancestrally marine, but as a result of numerous transitions across the salinity divide, freshwater species outnumber marine ones^7,12^. In addition to marine and freshwater specialists, many lineages include “euryhaline” generalists that survive a broad salinity range. Selection should favor this type of plasticity in species that experience environmental fluctuations^9,13,14^ such as the rapid salinity changes that occur in coastal and estuarine biomes^15^. The ability of populations to survive abrupt environmental change rests on their ability to survive the initial cellular stress. Certain stress responses are widely conserved across species^16–18^, but lineage-specific regulation of stress responses have also been identified^19^ and are important because species able to mount more robust stress responses may be more likely to successfully colonize new environments. Through controlled RNA-seq experiments built upon decades of laboratory studies^20–22^, the physiological responses by diatoms to low salinity are coming into focus^11,23–25^, but comparative studies are necessary to show how some lineages have established a foothold in inland waters while others have not.

To better understand mechanisms of marine–freshwater transitions, we used experimental RNA-seq to characterize the short-term, minutes-to-hours, response to hyposaline stress in the diatom, *Skeletonema marinoi*. We compared this with published data^23^ on the acclimated state of *S. marinoi* exposed to weeks of hyposaline conditions to develop a more complete temporal model of salinity acclimation. We then compared the response of *S. marinoi* to that of another diatom, *Cyclotella cryptica*. The two species share a common (marine) ancestor 90 million years ago (Fig. 1). *Skeletonema marinoi* has deep marine ancestry with a recently evolved, modest tolerance to low salinity, growing in habitats generally ranging from marine to brackish^26,27^ (Fig. 1). *Cyclotella cryptica* is a more robust generalist that grows in salinities ranging from marine to freshwater^24^. It is part of a clade with deep freshwater ancestry and repeated traversals across the salinity gradient that gave rise to marine specialists, freshwater specialists, and salinity generalists (Fig. 1). The divergent ecologies and phylogenetic histories of *S. marinoi* and *C. cryptica* combined to offer novel mechanistic insights into the mitigation of hyposaline stress in fluctuating environments and, more broadly, clues about the properties of successful freshwater colonists.

**Fig. 1.**
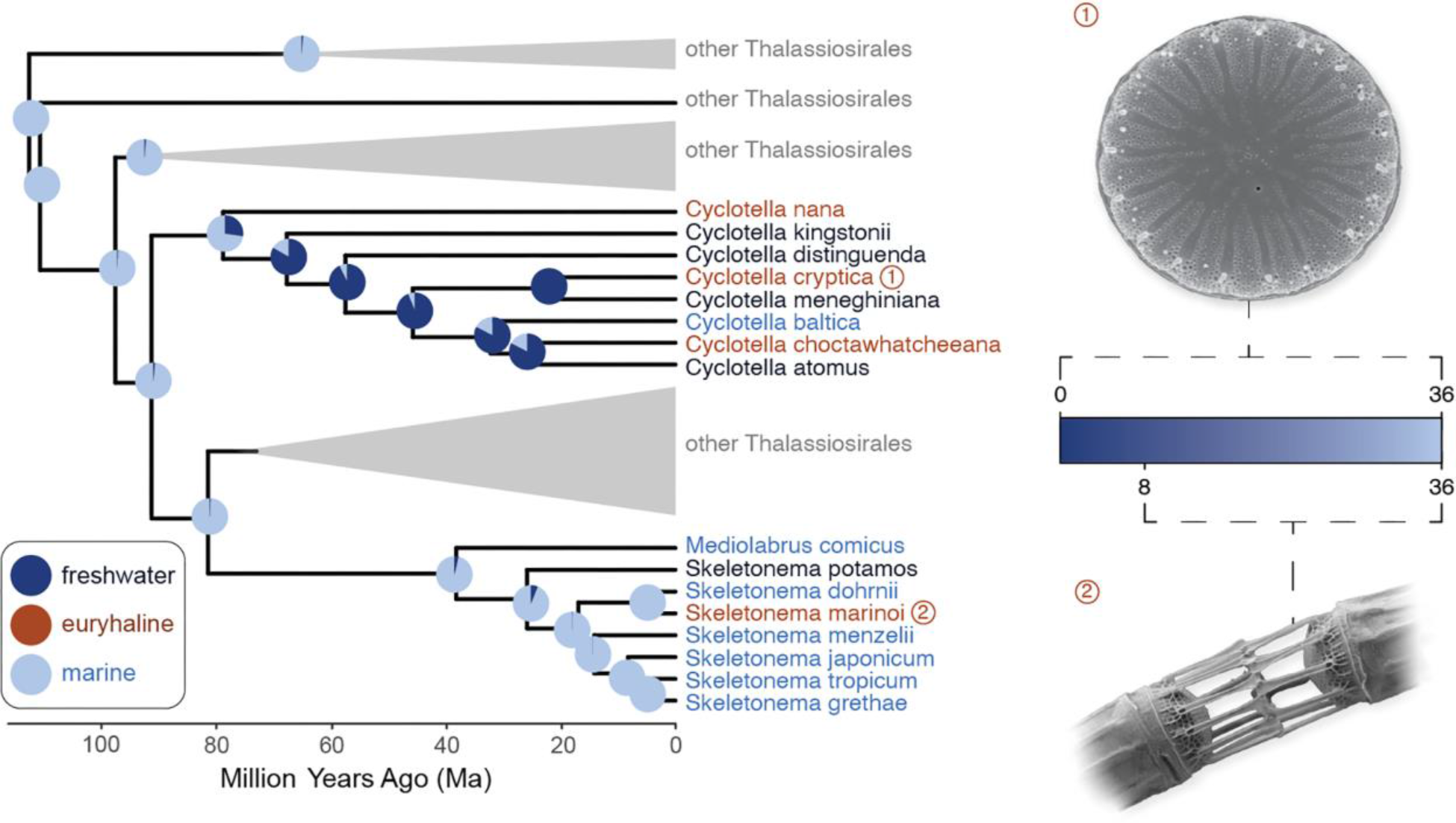
The distinct evolutionary trajectories of two euryhaline diatoms, *Skeletonema marinoi* and *Cyclotella cryptica*, following their split from a marine ancestor. *Cyclotella cryptica* is embedded inside a clade with deep freshwater ancestry and several marine–freshwater transitions. *Skeletonema marinoi* has deeper marine ancestry with fewer subsequent transitions. Approximate salinity tolerance is shown by the gradient ranging from freshwater (blue) to marine (white). Pie charts represent the probability of habitat (marine or freshwater) at ancestral nodes. Figure adapted from^12^.

## RESULTS & DISCUSSION

### Hyposalinity stress induces substantial remodeling of the transcriptome

We exposed *S. marinoi* strain CCMP3694 to hyposalinity stress by transferring cells from their native salinity at 24 grams salt per liter (ASW 24)^23^ to low salinity (ASW 8), and sequenced the transcriptome at seven timepoints ranging minutes to hours post transfer. Both the control and stressed cells experienced an initial lag within the first two hours with little or no growth in the control and some mortality in the treatment (Suppl. Fig. S1). The control increased growth from 2 h onwards, whereas stressed cells resumed growth at 4 h, albeit at a lower rate than the control (Suppl. Fig. 1). By 7 days, the control and treatment cultures had accumulated similar biomass.

Hyposalinity stress caused profound remodeling of the transcriptome. Of the 22,440 genes in *S. marinoi*’s genome, 14,860 were differentially expressed in at least one time point. The peak response, measured by the number of differentially expressed genes, occurred 2 h following stress exposure (8,086 genes) (Fig. 2a), coinciding with resumption of growth (Suppl. Fig. S1). The largest number of expression changes occurred 1–4 hours post-treatment (Fig. 2a). This was supported by multidimensional scaling of the top 500 differentially expressed genes, where the 1–4 h time points were most distant from the control (Fig. 2b). Notably, this time window corresponded with the initial decrease in biomass (1–2 h) and resumption of growth (2–4 h) (Suppl. Fig. S1). In contrast, the fewest differentially expressed genes were measured at 8 h, which was most closely aligned to the control in ordination space (Fig. 2b), indicating that expression profiles at the beginning and end of the experiment were more similar to each other than to intermediate time points (Fig. 2b). By 8 h *S. marinoi* had fully resumed growth and gene expression was returning to baseline levels (Figs 2a, Suppl. Fig. S1).

**Fig. 2.**
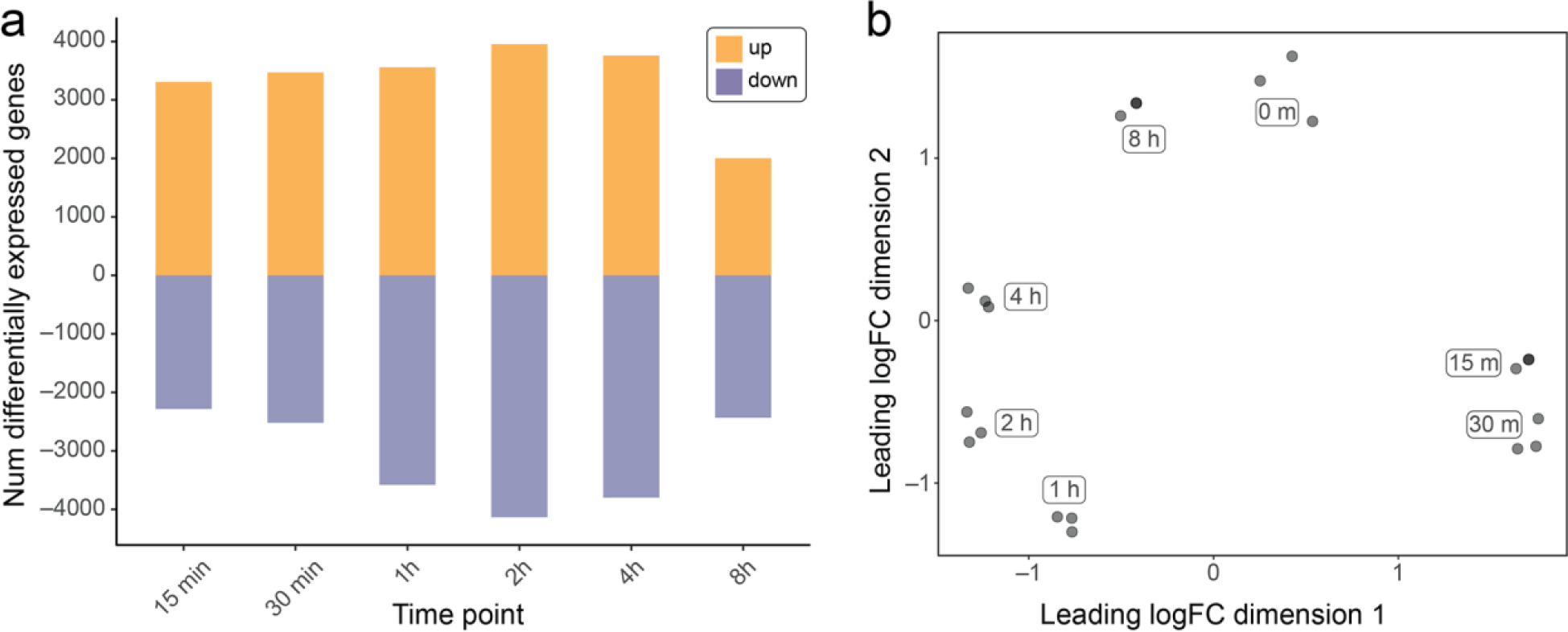
Hyposaline stress remodels the transcriptome of *Skeletonema marinoi*. a. Number differentially expressed at each time point following exposure to low salinity. The number of significantly upregulated genes are shown in purple, and significantly downregulated genes are shown in green. **b.** Multidimensional scaling plot, showing distinct patterns of gene expression at each time point, based on logFC changes in the top 500 differentially expressed genes.

Genes most critical to the stress response should have sustained patterns of differential expression across time^11^, so we focused on the 10,050 genes differentially expressed at two or more consecutive time points. Hierarchical clustering of these genes revealed seven groups defined by distinct temporal dynamics of gene expression (Fig. 3). Six of seven groups showed the largest magnitude of expression responses in the 1–4 h time period, confirming the peak stress response identified by the total number of differentially expressed genes (Fig. 3b).

**Fig. 3.**
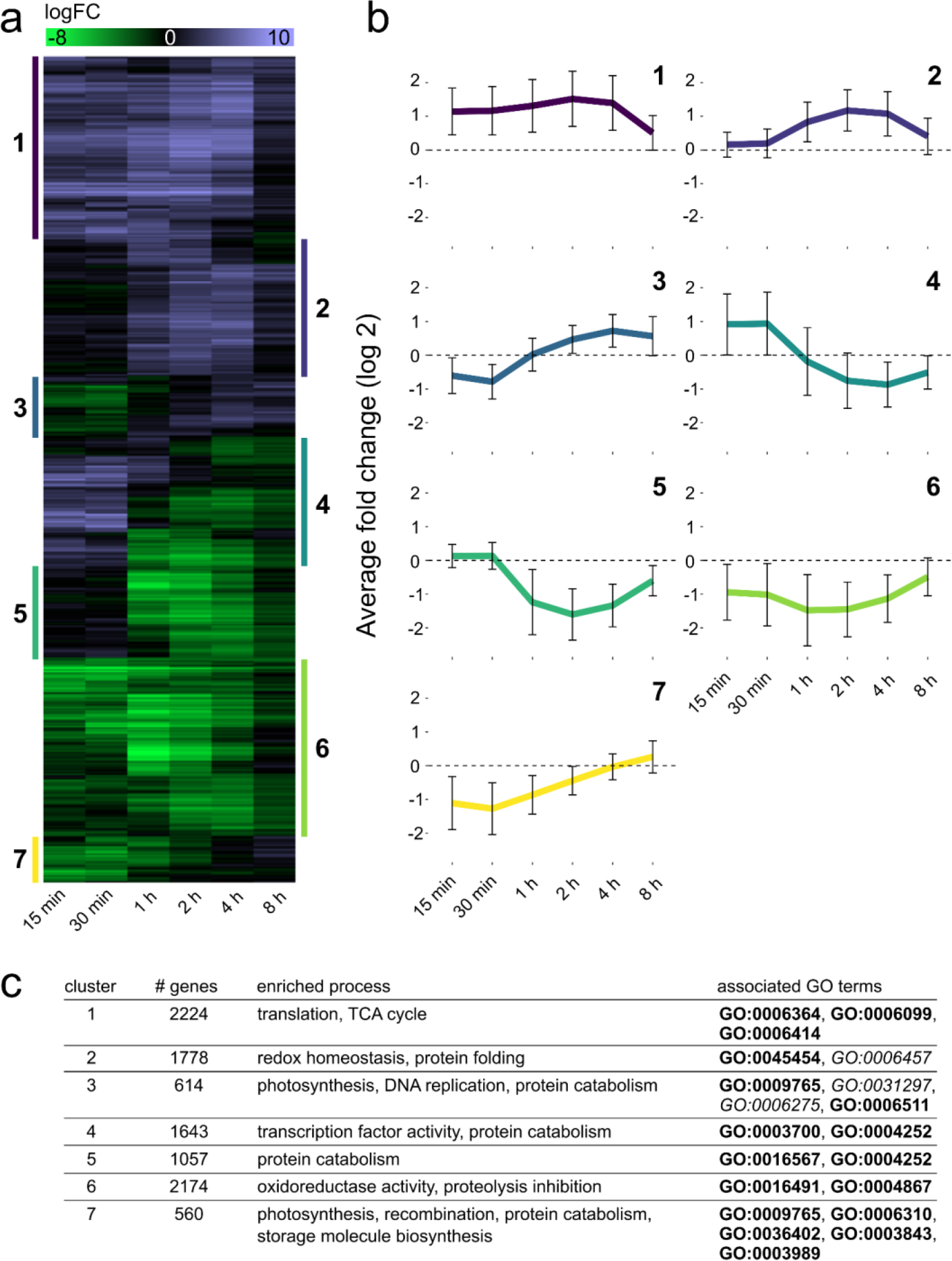
Differentially expressed genes exhibit distinct temporal dynamics during the hyposalinity stress response. a. Heatmap of 10,050 genes (Y axis) differentially expressed in ≥ 2 consecutive time points (X axis), sorted by similarity in gene expression across time points. Genes were classified into seven clusters based on shared patterns of gene expression. **b.** Average expression of genes assigned to each cluster. Error bars indicate ± one standard deviation. Cluster numbers are indicated on the sides of the heatmap in panel A. **c.** Number of genes, enriched biological processes, and specific GO terms for each cluster. GO terms in bold and italic indicate enrichment at p-value < 0.001 and < 0.01, respectively. The full list of significant GO terms is available on Zenodo.

### Distinct temporal dynamics in expression patterns during hyposalinity stress

The abrupt shift to low salinity induced an immediate transcriptomic response in *S. marinoi* (Fig. 3). Within the first hours of hyposaline stress, *S. marinoi* maintained homeostasis through a multiphased stress response, culminating in the onset of acclimation at approximately 8 h (Fig. 3). Below, we highlight distinct phases of this response.

### Sustained increase in protein biosynthesis

*Skeletonema marinoi* increased protein expression throughout the 8 h, reflecting altered metabolism and/or replenishment of stress-damaged proteins. Specifically, most tRNA synthetase genes were upregulated across the time series (Suppl. Fig. S2, p ≈ 4.2e-7, Fisher’s exact test), and 86 ribosomal proteins were upregulated at all time points (p ≈ 0, Fisher’s exact test). Genes involved in nitrogen metabolism were also mostly upregulated throughout the experiment (Suppl. Fig. S3), including ones involved in the transport and assimilation of ammonium, urea, and nitrate/nitrite. This pattern, alongside upregulation of phosphate transporters in early time points, is consistent with elevated nutrient demands associated with increased protein biosynthesis.

### Stress mitigation phase (15–30 min)

During the first 30 min of hyposalinity stress, *S. marinoi* experienced a reduced growth rate and a marked shift in energy metabolism. Growth halted immediately (Suppl. Fig. S1), which was reflected in the downregulation of DNA replication/recombination. Downregulation of chlorophyll biosynthesis, light-harvesting proteins, Calvin cycle genes, gluconeogenesis, and storage molecule biosynthesis (fatty acids and chrysolaminarin) suggests overall downregulation of photosynthesis and energy storage immediately upon stress exposure (Fig. 3b-c, Suppl. Figs S4–S7). Instead, *S. marinoi* upregulated chrysolaminarin degradation, the last irreversible step of glycolysis (pyruvate kinase), and the TCA cycle (Fig. 3b-c, Suppl. Fig. S4). Together, these patterns suggest that stress mitigation hinges on a metabolic shift characterized by rapid energy production through utilization of storage molecules and metabolic intermediates from glycolysis and the TCA cycle.

Transcriptional changes indicated that *S. marinoi* experienced acute stress during the first 30 minutes. This was evidenced by upregulation of: (i) heat shock proteins (Suppl. Fig. S8), which are molecular chaperones that direct damaged or misfolded proteins to proteases^43,44^, (ii) genes involved in mitotic DNA damage and integrity checkpoint signaling, which prevents mitosis in the presence of DNA damage^45^, and (iii) serine-type endopeptidases, which are likely involved in the degradation of stress-damaged proteins^46^. Many heat shock proteins were subsequently downregulated after 1 hour, suggesting that the acute stress was largely mitigated by that point (Suppl. Fig. S8). During the stress mitigation phase, *S. marinoi* also induced genes that mitigate the effects of reactive oxygen species (ROS), which has been found in other algae^47,48^. ROS-mitigating processes included upregulation of: (i) violaxanthin de-epoxidase, involved in the energy-dissipating xanthophyll cycle^49^, (ii) biosynthesis of biliverdin, a scavenger of oxygen radicals^50^, (iii) biosynthesis of polyamines that function in ROS management^51^ and osmotic balance^52^, and (iv) ROS scavengers, including superoxide dismutase, which is a first line of defense against ROS in plants and algae^53,54^ (Suppl. Figs S7, S9-S10).

At the onset of stress, *S. marinoi* upregulated key pathways involved in osmotic stress mitigation (Suppl. Fig. S10). Osmolytes are low-molecular-weight molecules whose intracellular concentrations maintain osmotic balance^55–57^. During hyposaline stress, we expect decreased expression of osmolyte genes. Indeed, several osmolyte biosynthesis genes were downregulated and osmolyte degradation genes were upregulated (e.g., taurine dioxygenase) in early time points or, in some cases, across all time points (Suppl. Fig. S10). Many of the strongest expression responses occurred during the stress mitigation phase, including methyltransferases involved in dimethylsulfoniopropionate (DMSP) and glycine betaine biosynthesis (Suppl. Fig. S10). Although proline is a well-characterized osmolyte indiatoms, expression patterns of genes involved in proline metabolism were inconsistent (Suppl. Fig. S10), suggesting proline might not be a universal osmolyte in diatoms^25^. Diatoms also maintain osmotic balance by regulating intracellular ion concentrations. Here, 11 of the 26 differentially expressed Na^+^ and K^+^ transporters were part of clusters upregulated at 15–30 min (Suppl. Fig. S11). Remaining ion transporters were confined to clusters downregulated across the entire time series (Suppl. Fig. S11). In addition, five amino acid ABC transporters were upregulated during the stress mitigation phase only (Suppl. Fig S12), consistent with removal of osmolyte-functioning amino acids from the cytosol early on^57,58^.

### Transition (1 h) and recovery (2–4 h) phases

Acute stress mitigation gradually transitioned to recovery, as expression levels at 1 h showed patterns that were a mix between the preceding mitigation and later recovery phases. The latter represents the peak response, as it shows the largest number of differentially expressed genes and coincides with upregulation of cell cycle genes and growth resumption (Figs 2a, 3b). Although many genes involved in osmotic and oxidative stress were no longer differentially expressed during recovery, several transporter and osmolyte genes continued to be differentially expressed (Suppl. Figs S10-S12). Similarly, many cell-compartment-associated peroxiredoxins and thioredoxins remained upregulated, often at larger magnitudes, at 2–4 h. Most cellular processes that were predominantly upregulated throughout the experiment reached peak upregulation during the recovery phase, most notably protein biosynthesis, chrysolaminarin degradation, and the TCA cycle (Suppl Figs S2, S4). In parallel, key irreversible steps of glycolysis were upregulated by 2 h (Suppl. Fig. S4), indicating that storage molecules continued to provide energy during the recovery phase. Genes involved in proteasome activity also became upregulated at 4 h, consistent with clearing of damaged or unnecessary proteins (Suppl. Fig. S13). Altogether, *S. marinoi* carefully balanced stress mitigation and energy production during the recovery phase.

### Pre-acclimation phase (8 h)

By 8 h, growth had resumed, several cellular processes that were initially downregulated became upregulated, and the overall magnitude of the gene expression responses declined (Figs 2, 3b), indicating that *S. marinoi* had begun to acclimate to low salinity^59^. This is confirmed by the large overlap in differentially expressed genes and pathways between the 8 h time point and a previous study of *S. marinoi* acclimated to ASW 8 for two weeks^23^ (Fig. 4). Acclimated and pre-acclimated cells increased storage molecule biosynthesis, suggesting *S. marinoi* had fully catabolized storage molecules for energy during the first hours of the response and were replenishing their stocks (Fig. 4, Suppl. Fig. S4). Similarly, although the TCA cycle was upregulated during our experiment, downregulation of the *bZIP14* transcription factor, which regulates the TCA cycle in diatoms^60^, from the recovery phase onward suggests the TCA cycle was becoming increasingly downregulated at 8 h and was, in fact, fully downregulated in acclimated cells at two weeks (Figs 3-4, Suppl. Fig. S4). This suggests that the TCA cycle plays an important role in supplying energy to the cell during acute stress but not acclimation.

**Fig. 4.**
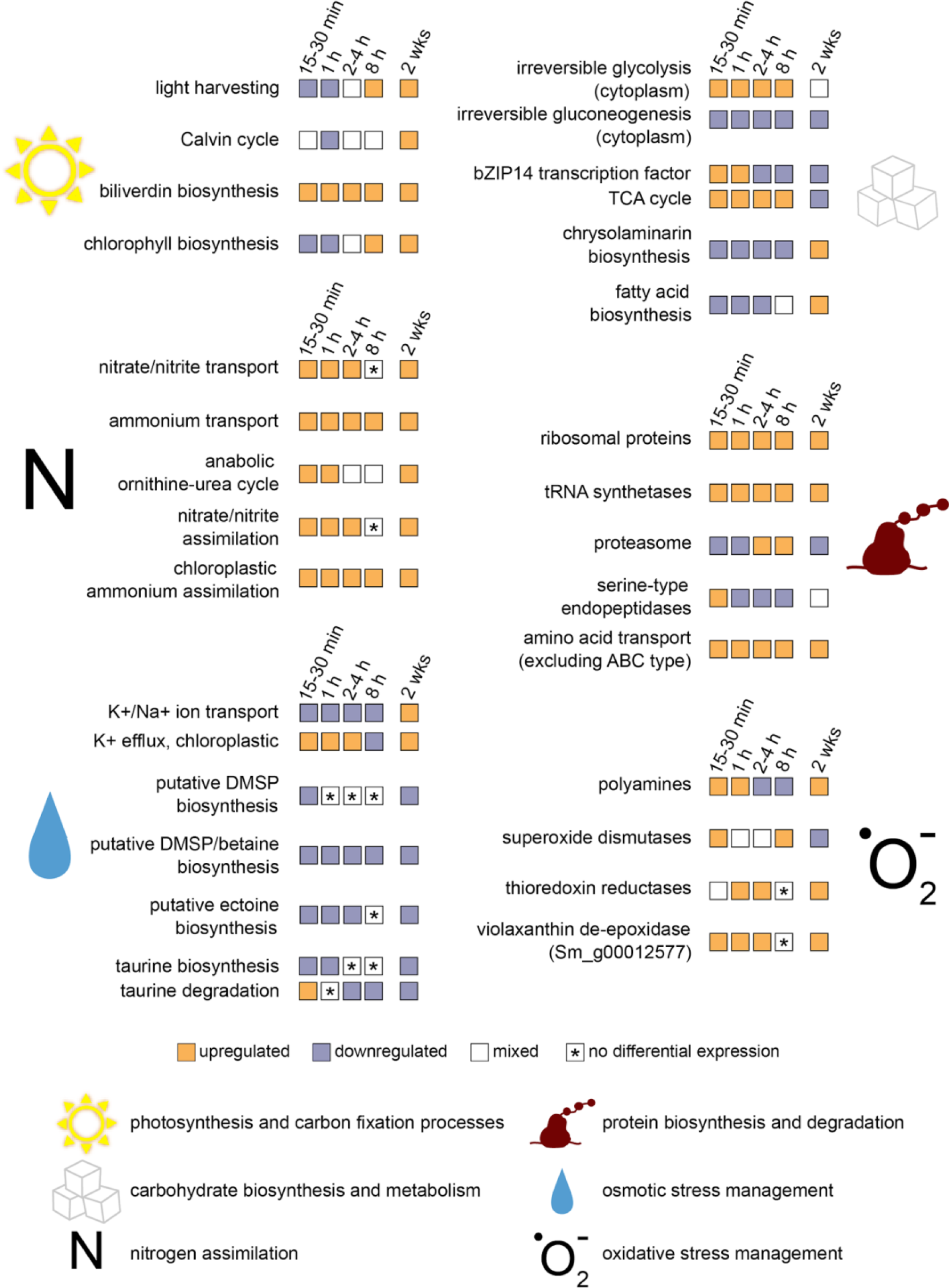
The hyposalinity stress response versus acclimation. Differentially expressed genes are involved in diverse cellular processes during acute stress (15 min to 8 h) and acclimation (2 weeks). The colored tiles represent the four phases of short-term expression: stress mitigation (15–30 min), transition (1 h), recovery (2–4 h), pre-acclimation (8 h), and acclimation (2 wks)^23^. Vertical spacing of tile rows indicates associations between processes (e.g., the *bZIP14* transcription factor regulates the TCA cycle). Tile color was determined based on the proportion of genes significantly up- or downregulated for that process. Purple: > 60% downregulated genes, Orange: tiles indicate > 60% upregulated genes, White: 40–60% up- and downregulated genes, Asterisks: no differentially expressed genes for that process.

Notably, several processes showed similar patterns during the early phases of the stress response and at two weeks of acclimation, but without differential expression, or expression in opposite directions, during the pre-acclimation phase (Fig. 4). These include the Calvin cycle, transport and assimilation of nitrate/nitrite, the ornithine-urea cycle, genes involved in the biosynthesis of osmolytes and polyamines, ion transporters, proteasome genes, thioredoxins, and violaxanthin-de-epoxidase (Fig. 4, Suppl. Figs S3, S6, S9-13). ROS-management strategies were generally downregulated or not differentially expressed during the pre-acclimation phase (except SOD) (Suppl. Fig. S9), but became upregulated at two weeks of acclimation, whereas the proteasome showed an opposite trend. This suggests that *S. marinoi* reached a new equilibrium for ROS management after full acclimation, namely one that prevented damage to cellular components and allowed for downregulation of proteasome activity. Overall, the contrasting expression patterns between the early time points in our experiment and long-term acclimation on the one hand, and the pre-acclimation phase on the other hand, suggest that the latter represents an ‘overcorrection’ for some pathways, as has been observed in macrobiota^61^. This might be caused by surplus metabolites, synthesized during the recovery phase, which triggers transient downregulation of their corresponding pathways by feedback inhibition until acclimated cells reach a new equilibrium. Similar feedback mechanisms have been observed for nitrogen metabolism and polyamine biosynthesis in bacteria and algae^62,63^.

### Conserved and diverged responses to hyposaline stress in two euryhaline diatoms

We compared the response to acute hyposalinity stress from *S. marinoi* with *C. cryptica* to reveal patterns of conservation and divergence^11^. Both experiments were completed simultaneously and used the same design, with the most notable difference being the magnitude of the hyposalinity exposure, which amounted to brackish conditions (ASW 8) for *S. marinoi* and freshwater (ASW 0) for *C. cryptica*. Both species underwent substantial transcriptional remodeling in the minutes and hours following hyposaline stress but eventually approached acclimation to decreased salinity. Taken together, this suggests that the two species experienced a comparable degree of stress despite differences in the salinity exposure. This is consistent with this strain of *S. marinoi* being a less robust euryhaline diatom, as it does not survive below approximately salinity 4.

### Conserved features of the hyposaline stress response

Several similarities in the transcriptional responses of the two species identify the conserved features of the response to hyposaline stress by diatoms. Both species experienced an immediate arrest in growth and widespread up- or downregulation of signal transduction kinases, including (i) histidine kinases, which are responsible for signal transduction across cell membranes and facilitate stress adaptation across the tree of life^64^, (ii) serine/threonine protein kinases, which are involved in stress responses in diatoms^65,66^, and (iii) cGMP-dependent protein kinases, which are important for salt stress responses in vascular plants^67^. Notably, these signal transduction kinases were targets of positive selection associated with adaptation of *S. marinoi* to the Baltic Sea environmental gradients, including low salinity^68^. Conserved features of the early phases of the stress response (15–30 min) include (i) oxidative stress management with ROS scavengers and polyamines, (ii) upregulation of heat shock proteins and associated transcription factors, (iii) downregulation of cell cycle genes and histones. In this early phase, both species also upregulated (i) plastid K^+^-efflux antiporters to counter osmotic pressure in the chloroplasts, (ii) phosphate and molybdate transporters, presumably to meet increased demands for resources allocated to damage repair and growth resumption, and (iii) a chitinase. *Cyclotella cryptica* forms B-chitin threads which might play a role in buoyancy adjustments under acute hyposalinity stress ^11^. However, given that *S. marinoi* does not form such chitin-structures, a general chitin response in both species suggests a broader role of chitin in hyposalinity stress mitigation, perhaps involving cell wall remodeling^11,69,70^. Many of the aforementioned processes were still upregulated at 2 h. However, both species now also upregulated cell cycle genes, indicative of resumed growth, whereas most heat shock proteins and associated transcription factors became downregulated. This pattern continued at 4 h, in addition to substantial upregulation of genes involved in protein translation. Finally, at 8 h, both species continued to upregulate translational activity and SOD.

### The swift, efficient, and orchestrated response to hyposaline stress in a diatom with freshwater ancestry

Despite many similarities, several important differences in the responses to hyposalinity stress between the two species highlighted key features that impart greater overall salinity tolerance in *C. cryptica*. Although *S. marinoi* experienced a milder salinity shift than *C. cryptica*, *S. marinoi* nevertheless had to mount a much stronger response in terms of both the number and magnitude of differentially expressed genes. Specifically,

*S. marinoi* differentially expressed a larger fraction of its genes (14,860 genes; 66%) during the time series than *C. cryptica* (10,566; 50%), including at each individual time point. Few enriched GO terms were shared between species at the same time points, and many that were shared were expressed in opposite directions, pointing to fundamentally different responses in the two species (Fig. 5). Notably, the majority of these opposite expression patterns were confined to 30–120 min, whereas shared GO terms at 4–8 h tended to be expressed in the same direction in both species, suggesting the responses of both species most strongly diverged during initial acute stress. Considering genes differentially expressed in both species, the magnitude of the response, expressed as logFC, was significantly greater in *S. marinoi* at nearly all time points (Fig. 6, Suppl. Fig. S14). Finally, the multiphased response of *S. marinoi*, spread over 8 h, was much quicker in *C. cryptica*, which mounted a stronger immediate response with peak gene expression during the initial stress mitigation phase (30–60 min), with gene expression returning to baseline levels by 4 h for many processes. By contrast, peak gene expression occurred during the recovery phase (2–4 h) in *S. marinoi*, suggesting *C. cryptica* directs the most effort towards immediate stress mitigation, whereas *S. marinoi* invests the most energy in its recovery.

**Fig. 5.**
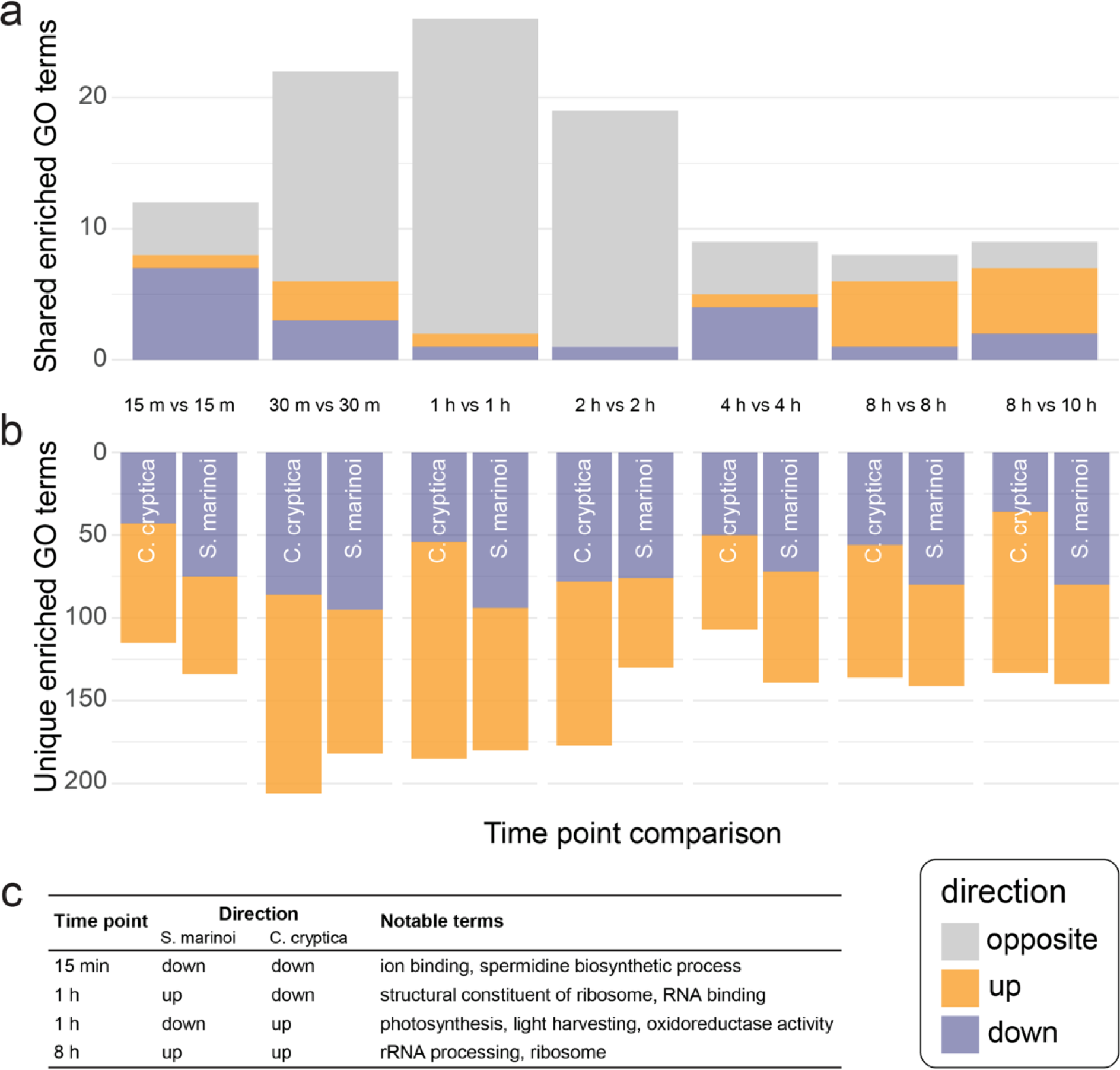
The distinct responses of two salinity generalists, *Skeletonema marinoi* and *Cyclotella cryptica*, to hyposaline stress. **a.** Number of enriched GO terms (p-value < 0.05) shared between the two species in the minutes and hours following exposure to hyposaline stress. **b.** Number of enriched GO terms (p-value < 0.05) unique to each species. **c.** Shared enriched GO terms with the greatest number of shared GO terms. A full list of enriched GO terms by category is available in our Zenodo repository.

**Fig. 6.**
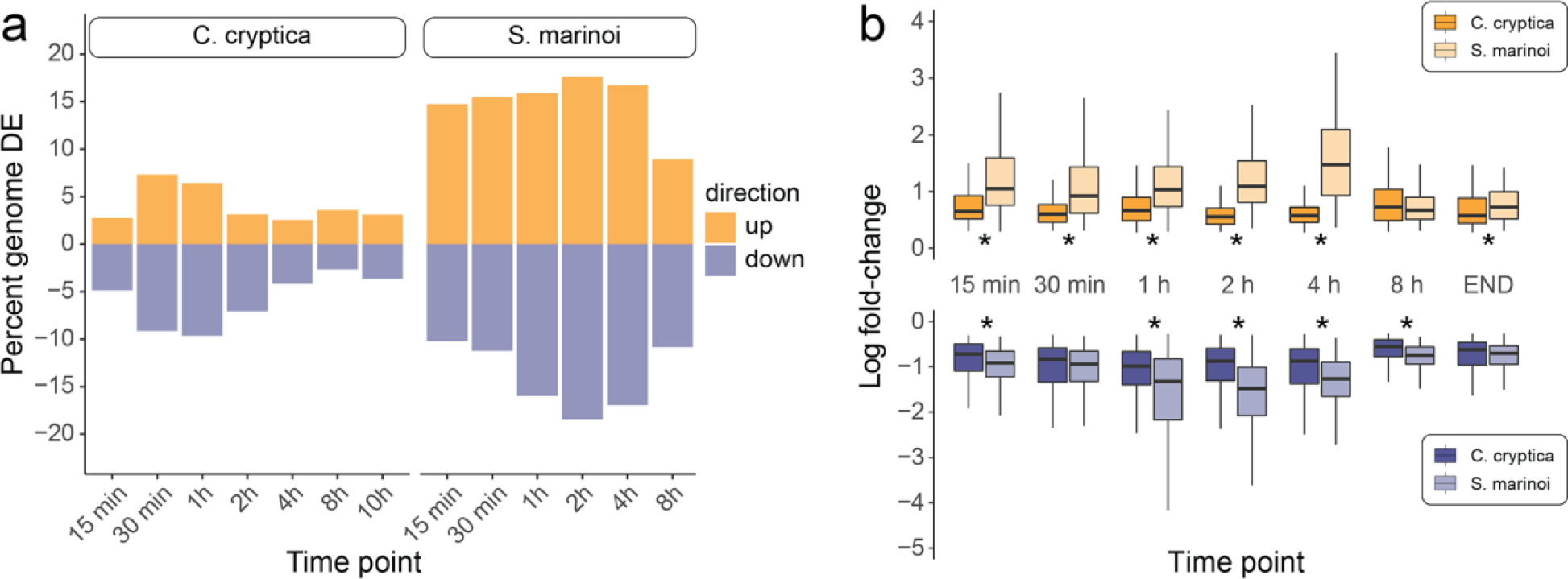
Differences in the strength of the hyposalinity stress response. Despite milder stress exposure, *S. marinoi* expresses more genes (**a**) at greater magnitudes (**b**) than *C. cryptica*. Values in (**b**) show the distribution of LFC values for shared homologs with significant differential expression in both species. Upregulated genes are on the top panel, downregulated on the bottom. If (in-)paralogs were present, their LFC was included individually. Stars indicate a significant difference (p-value < 0.05) in population mean ranks by a two-sided, two-sample Wilcoxon rank sum test. Outliers not shown.

Major differences between species fell into two categories: (i) activity of osmolytes and ion channels, and (ii) management of oxidative stress. *Cyclotella cryptica* restores osmotic balance during acute hypo-salinity stress by transporting K^+^ into the cytosol, and Na^+^ and H^+^ out of the cytosol, whereas osmolytes play only a minor role (e.g., biosynthesis of DMSP and glycine-betaine were not differentially expressed). *Cyclotella cryptica* upregulates 32 K^+^ or Na^+^ transporters across multiple time points, and regulates an additional 66 K^+^ or Na^+^ transporters at single time points with roughly equal distribution of up- and downregulation across transporters, consistent with subfunctionalization of paralogs. *Skeletonema marinoi*, by contrast, regulates osmotic stress by strongly reducing osmolyte levels, and much less so through ion transport. In *S. marinoi* fewer (n = 26) K^+^ or Na^+^ transporters were differentially expressed at consecutive time points, and most of these (n = 17) were primarily downregulated (Suppl. Fig S11). Moreover, only 11 ion transporters were differentially expressed at single time points, with near equal distribution of up- and downregulation. Strong upregulation of amino acid transporters in *S. marinoi*, including five ABC transporter-binding proteins at 15–30 min, further suggests a dominant role for osmolytes in osmoregulation by *S. marinoi*, as we expect these transporters to actively expel excess osmolytes from the cell. *Cyclotella cryptica* either downregulated or did not differentially express most of its amino acid transporters throughout the time series.

The canonical response to environmental stress in many organisms involves transient repression of ribosome biogenesis and upregulation of stress defense genes^71,72^. This was also observed in the response to hyposaline stress by *C. cryptica* ^11^, but *S. marinoi* instead upregulated both translation and protein degradation across the entire time series. This suggests that whereas a reduction in translation reflects the growth arrest in *C. cryptica*, sustained and energy-demanding degradation and replenishment of ROS-damaged proteins drove increased protein degradation and translation in *S. marinoi* for a full 8 h. This suggests that protein damage was more extensive and prolonged in *S. marinoi* compared to *C. cryptica*. Furthermore, most light-harvesting proteins were upregulated in *C. cryptica*, whereas more than half were downregulated in *S. marinoi* (Suppl. Fig. S5). One possibility is that indirect quenching of overexcited chlorophyll prevented ROS generation in *C. cryptica*^73^, whereas *S. marinoi* limited ROS generation by decreasing photosynthesis altogether (Fig. 3b-c), which is a strategy used by some plants^74^. These observations suggest that the oxidative stress induced by low salinity in *S. marinoi* may be too severe to be managed by increased photoprotection mechanisms alone.

The long-term acclimation strategies also differed substantially between *S. marinoi* and *C. cryptica*. For *C. cryptica*, the acclimated state was highly distinct from the short-term stress response, as most genes and pathways differentially expressed during short-term stress were not so in acclimated cells, and for those genes that were, most were expressed in opposite directions in stressed versus acclimated cells^11^. In *S. marinoi*, by contrast, many genes and processes differentially expressed under short-term stress remained so in acclimated cells (Fig. 4). This indicates that acclimated *C. cryptica* reaches a new homeostasis with limited energy requirements, whereas *S. marinoi* mounts a prolonged stress response to continue growth in hyposaline conditions. It is worth noting here that the length of acclimation differed between the two species, so we cannot rule out that some of the differences reflect further changes in the acclimated state between two weeks (*S. marinoi*) and three months (*C. cryptica*).

## CONCLUSIONS

Using time-resolved transcriptomics, we found fundamental differences in the acute short-term stress responses and long-term acclimation strategies of two euryhaline diatoms exposed to low salinity. Despite considerably milder exposure, the weaker of the two generalists, *S. marinoi*, mounted a stronger and more prolonged response to hyposaline conditions than *C. cryptica*. In cases where successful stress management requires higher levels of gene expression, like in *S. marinoi*, its physiological limits and maximum energy demands will be reached at a lower dose of environmental stress, allowing for less tolerance to environmental extremes than species that can survive with a smaller and less energy-demanding response. This was evident in the divergent strategies for mitigating oxidative and osmotic stress, which offer important clues about the comparatively broader salinity tolerance of *C. cryptica*. The greater efficiency of ROS management in *C. cryptica* appears to have reduced the period of acute stress, allowing it to recover and resume growth sooner than *S. marinoi*. In addition, we hypothesize that the regulation of osmotic pressure through increased ion transport in *C. cryptica* is faster and more efficient than the osmolyte- dominated response of *S. marinoi*. Finally, the distinct gene expression profile of fully acclimated *C. cryptica* cells^11^ suggests it is better able to settle in comfortably to lower salinities than *S. marinoi*, where the expression profiles of acclimated cells more closely resemble stressed cells. Taken together, our data suggest that early and efficient responses to oxidative and osmotic stress, together with a tailored acclimation state, confer overall broader salinity tolerance.

Marine–freshwater transitions have occurred many times and in both directions throughout diatom evolution^7,75^, including in *Cyclotella*, which includes salinity generalists as well as marine and freshwater specialists (Fig. 1). Ancestral state reconstructions highlight a deep freshwater colonization event, followed by tens of millions of years of freshwater ancestry that subsequently gave rise to marine specialists and salinity generalists (Fig. 1)^12,75^. We hypothesize that these repeated transitions are examples of adaptive phenotypic plasticity^9^, in which the cellular mechanisms underlying the broad plasticity of *C. cryptica* are the same ones that allowed *Cyclotella* to become established in freshwaters originally. Long-term retention of key mechanisms—the ones distinguishing *C. cryptica* from *S. marinoi*—have allowed *Cyclotella* to subsequently, and repeatedly, go on to specialize in marine or freshwaters (Fig. 1) through genetic assimilation^9,76^. The longer marine ancestry of *S. marinoi* suggests its strategies to manage hyposaline stress are more recently evolved, less refined, and less plastic. Our experimental strain of *S. marinoi* originated from the marine North Sea, but strains locally adapted to low-salinity reaches of the Baltic Sea^26,77^ might be able to mount stress responses more resemblant of *C. cryptica*. Indeed, *S. marinoi* exhibits both genomic and transcriptional variation associated with the salinity gradient across the Baltic Sea^23,68^.

These results suggest that similarities in environmental stress responses across species are likely limited to shared ancestral mechanisms constituting part of a core stress response, whereas lineage-specific aspects may better predict survival to environmental perturbations on short timescales, and successful colonization of new habitats on longer timescales. These questions have taken on increased urgency in the context of climate change, where evolutionary history will play a role in determining which lineages survive and adapt to changing ocean conditions^78^.

## MATERIALS & METHODS

### Sample collection and experimental design

*Skeletonema marinoi* strain CCMP3694 was germinated from a resting cell collected in the North Sea near Gothenburg, Sweden (sampling year: 2014, germination year: 2017), and grown at 15 ℃ and 21.5 μmol photons⋅m^-2^⋅s^-1^ irradiance under a 16:8 light:dark cycle. Cells were maintained in artificial seawater with 24 grams salt per liter (ASW 24), the native salinity of the strain^23^.

*Skeletonema marinoi* cannot survive freshwaters, so hyposaline stress was induced by transferring cells from ASW 24 to ASW 8. Cells were inoculated into three 1 L flasks with ASW 24 prior to the experiment and growth was monitored with a Fluid Imaging Technologies Benchtop B3 Series FlowCAM particle counter. Upon reaching exponential growth, cells were enumerated with the FlowCam, concentrated by centrifugation (800 rcf, 3 min), and inoculated into 50 mL Falcon tubes containing ASW 24 (control) or ASW 8 (treatment), resulting in 3×10^6^ cells/tube (40 mL). Cells were collected for RNA-seq at seven time points: 0 min, 15 min, 30 min, 1 h, 2 h, 4 h, and 8 h. The control was collected alongside the 0 min treatment, immediately after inoculation at the start of the experiment. The remaining tubes were held at 15 ℃ under constant illumination (20 μmol photons⋅m^-2^⋅s^-1^) and gentle agitation with a Boekel Scientific wave rocker until collection. At each time point, cell pellets were concentrated by centrifugation (400 rcf, 3 min), flash-frozen in liquid nitrogen, and stored at –80 ℃. The experiment was performed in triplicate, resulting in 24 RNA-seq samples.

### Sequencing and read processing

We extracted RNA and prepared Illumina libraries in four batches. To minimize batch effects, samples were randomized using the “sample” function in R v4.05 (R Core Team, 2020) prior to both RNA extraction and library construction. RNA was extracted with a QIAGEN RNeasy Plant Mini Kit and sequencing libraries were prepared with a KAPA mRNA HyperPrep kit. Indexed libraries were multiplexed and sequenced on a single Illumina HiSeq 4000 lane (paired-end, 100 bp).

A total of 696,475,182 reads were sequenced. Reads were trimmed with kTrim v1.1.0 (parameters: -t 15 -m 0.5)^28^ and mapped to the *S. marinoi* reference genome v1.1 using STAR v2.7.3a^29^ with default settings and intron sizes ‘--alignIntronMin 4’ and ‘--alignIntronMax 17105’. Read counts were estimated using HTSeq v0.11.3 in *union* mode^30^. Gene annotations and protein localization predictions were obtained from Pinseel et al.^23^.

### Differential expression analysis

We analyzed transcript counts in R v4.0.2. Only genes with at least 1 count per million (CPM) in >3 samples were retained. We used edgeR v3.30.3 to adjust for variation in library size and composition using the trimmed mean of M-values (TMM) method and to fit a quasi-negative binomial general linear model (GLM) for each gene^31,32^. We used stageR v1.10.0^33^ to identify differentially expressed genes in each contrast between an experimental timepoint and time 0 (t=0 min) at ASW 8 with a false discovery rate (FDR) of 1%^33,34^. Genome-wide differences in gene expression among time points were visualized with multidimensional scaling using limma v3.44.3^35^, based on the top 500 genes with the greatest log2-fold changes between each pair of samples. Cluster v3.0 and Java Treeview were used to sort genes with similar expression patterns into seven manually delimited clusters^36,37^. We performed Gene Ontology (GO) term enrichment in topGO v2.40.0^38^ using the *elim* algorithm and Fisher’s exact test to identify functional similarities within clusters and time points. All GO terms identified in the genome of *S. marinoi* were used as the background set. GO terms with p-value <0.05 were considered significant, and redundant GO terms were removed using REVIGO (accessed on 22 November 2021) with a similarity cutoff of 0.5 and the SimRel score as similarity measure^39^.

### Comparison of the hypo-salinity stress responses of two euryhaline diatoms

A major goal of this study was to characterize conserved and divergent features of the short-term response to hyposaline stress in diatoms. To this end, we compared the responses of two diatoms, *S. marinoi* (this study) and *C. cryptica*^11^ (Fig. 1). Both experiments were carried out simultaneously in the same lab and using the same bioinformatic workflow, allowing direct comparisons. Differences between the two experiments included: (1) the duration—an additional time point at 10 h was taken for *C. cryptica*, and (2) the magnitude of the hypo-salinity shock—*S. marinoi* has a lower salinity tolerance (lower boundary is 2.5 grams salt per liter, though many strains do not survive below 5–8) than *C. cryptica* (lower boundary is 0)^22,40,41^, so *S. marinoi* was transferred from ASW 24 to ASW 8 whereas *C. cryptica* was transferred from ASW 32 to ASW 0.

To compare the responses of the two species, we clustered the predicted proteins from *S. marinoi* and *C. cryptica* with OrthoFinder v.2.2.6 into orthogroups^42^. Orthogroups can contain a mix of orthologs and paralogs, collectively referred to here as homologs. Differences in expression levels between shared homologs were assessed using two-sided two-sample Wilcoxon tests in base R’s *wilcox.test()* function with a significance cutoff of 0.05. We first tested for differences in raw expression levels using an unpaired Wilcoxon rank sum test, evaluating the distribution of LFC values for all significantly expressed homologs by rank, rather than absolute expression, to control for differences in stress treatments between species. Second, assuming the overall physiological response is determined by cumulative expression of all homologs at a given time point, a second test used the summed expression values of all homologs in each species, resulting in one expression value per orthogroup per species at each time point, which were compared with a Wilcoxon signed rank test (paired data). Since we did not have a 10 h time point for *S. marinoi*, we compared the 8 h time point of *S. marinoi* with *C. cryptica* at the 8 h and 10 h end points of that experiment.

## AUTHOR CONTRIBUTIONS

KJJ, KMD, AJA, and JAL conceived and designed the study. KJJ and KMD conducted the experiments. KJJ performed data analysis with support from EP and KMD. KJJ, EP, and AJA wrote the manuscript. AJA, EP, and JAL supervised the study. JAL edited the manuscript. All authors read and approved the final manuscript.

## DATA AVAILABILITY

The RNA-seq reads are available from the Sequence Read Archive (NCBI) under project number PRJNA1055154. The diatom strain used in this study is available from the Bigelow laboratory, National Center for Marine Algae and Microbiota under accession CCMP3694. The bioinformatics code and data files used/created in this study are available from our Zenodo repository (10.5281/zenodo.10999657).

## ACKNOWLEDGEMENTS

This study is based upon work supported by grants from the Simons Foundation (403249 to AJA and 725407 to EP), Science Foundation Flanders, FWO (1221323N to EP), National Science Foundation (DEB- 1651087 to AJA and MCB-1941824 to JAL), and multiple grants from the Arkansas Biosciences Institute. EP benefited from postdoctoral fellowships from Fulbright Belgium and the Belgian American Educational Foundation. This research used resources available through the Arkansas High Performance Computing Center, which is funded through multiple NSF grants and the Arkansas Economic Development Commission. We thank Anna Godhe for providing sediment samples from the North Sea and Wade Roberts for providing the phylogenetic tree. The co-first authors are listed in alphabetical order.

