## Supplemental files for "The divergent responses of salinity generalists to hyposaline stress provide insights into the colonization of freshwaters by diatoms"

### SUPPLEMENTARY FIGURES

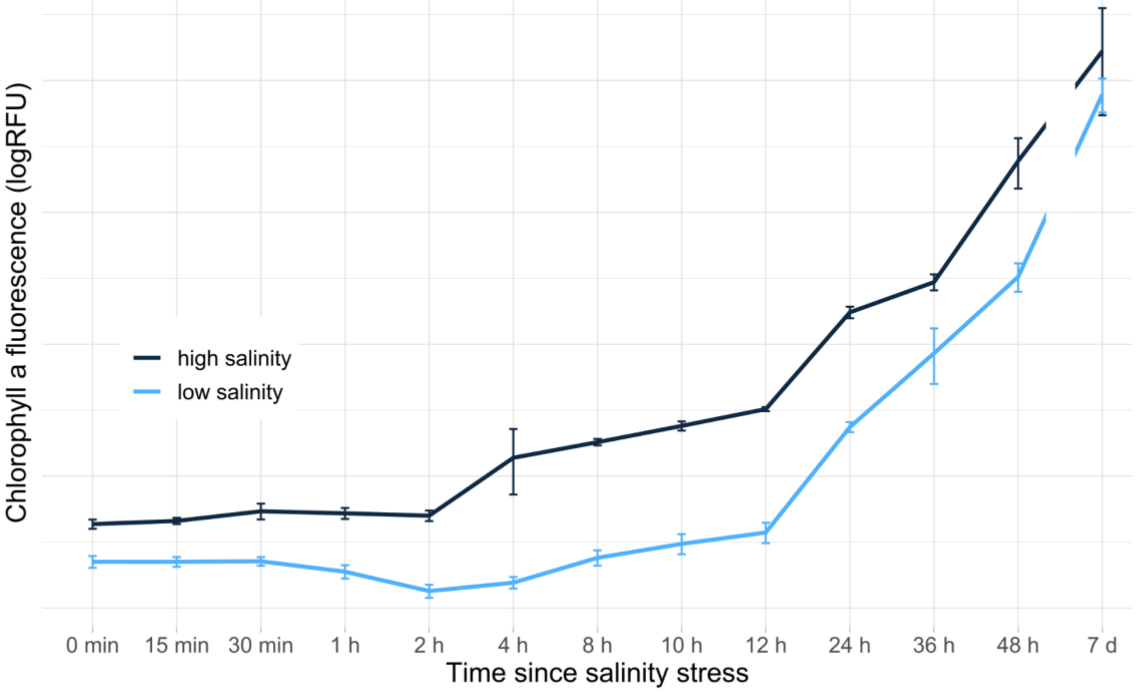


**Supplementary Fig. S1 | Growth of *S. marinoi* under acute hyposaline stress.** Chlorophyll *a* fluorescence (log2 relative fluorescence units [RFU]) as a proxy for growth in *S. marinoi* over 7 days. Growth is shown for high (24, control) and low (8) salinity. Error bars show standard deviations from biological triplicates.

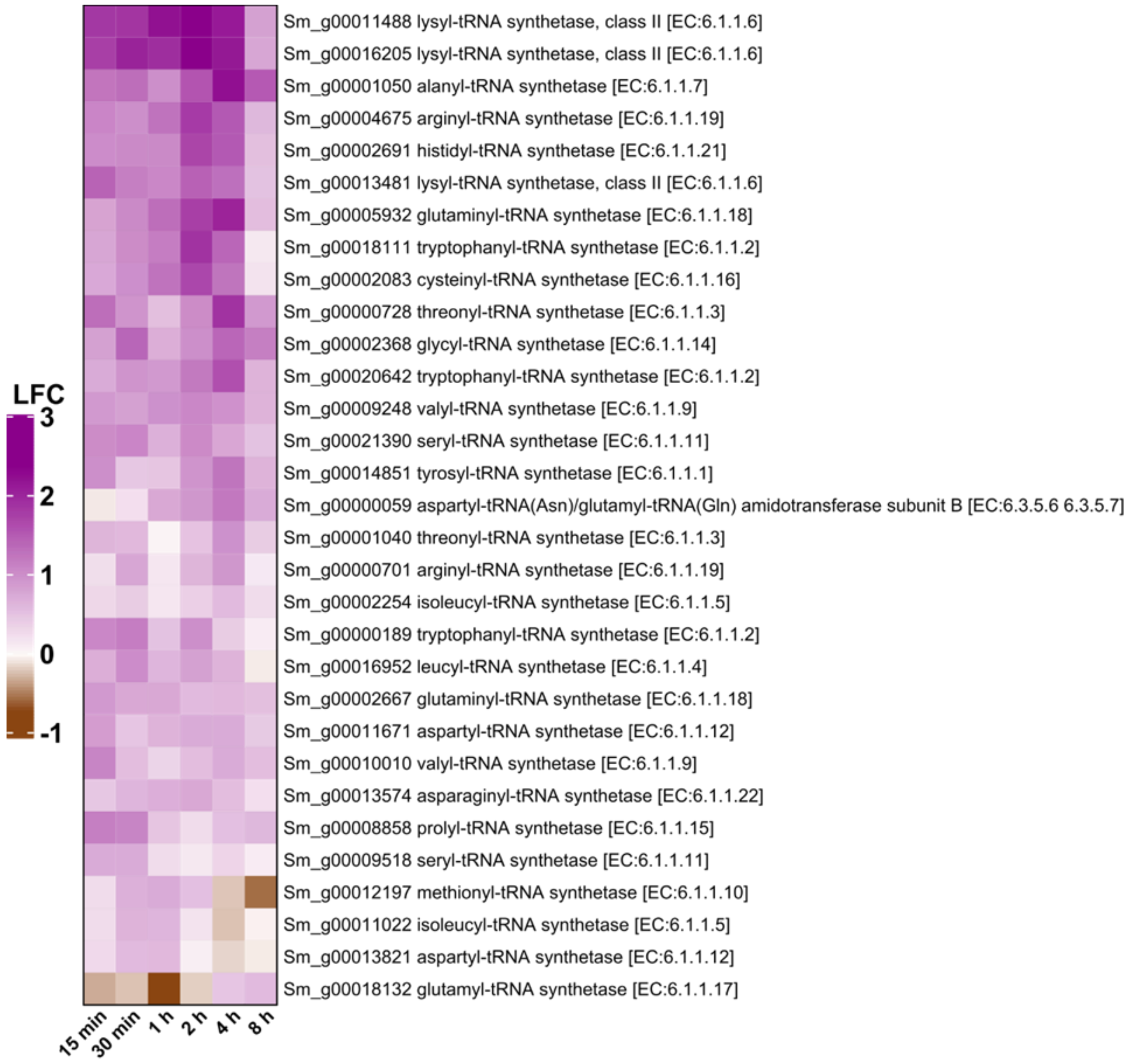


**Supplementary Fig. S2 | Time-resolved expression of tRNA biosynthesis.** Heatmap showing the expression of differentially expressed genes involved in tRNA biosynthesis over the 8-hour time series. All shown genes were significant in two or more consecutive time points.

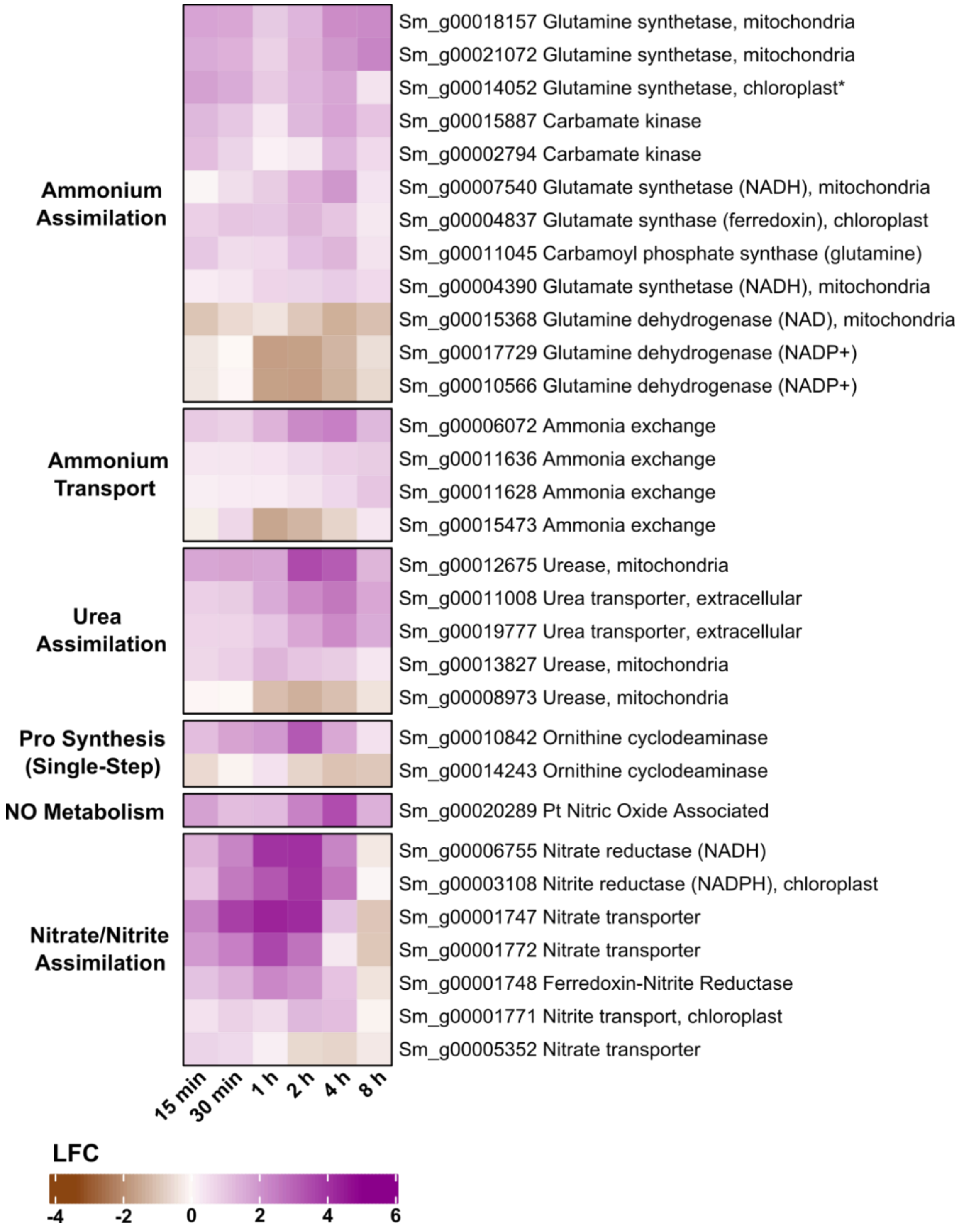


**Supplementary Fig. S3 | Time-resolved expression of nitrogen metabolism.** Heatmap showing the expression of differentially expressed genes associated with nitrogen assimilation over the 8-hour time series. All shown genes were significant in two or more consecutive time points. Continued on the next page.

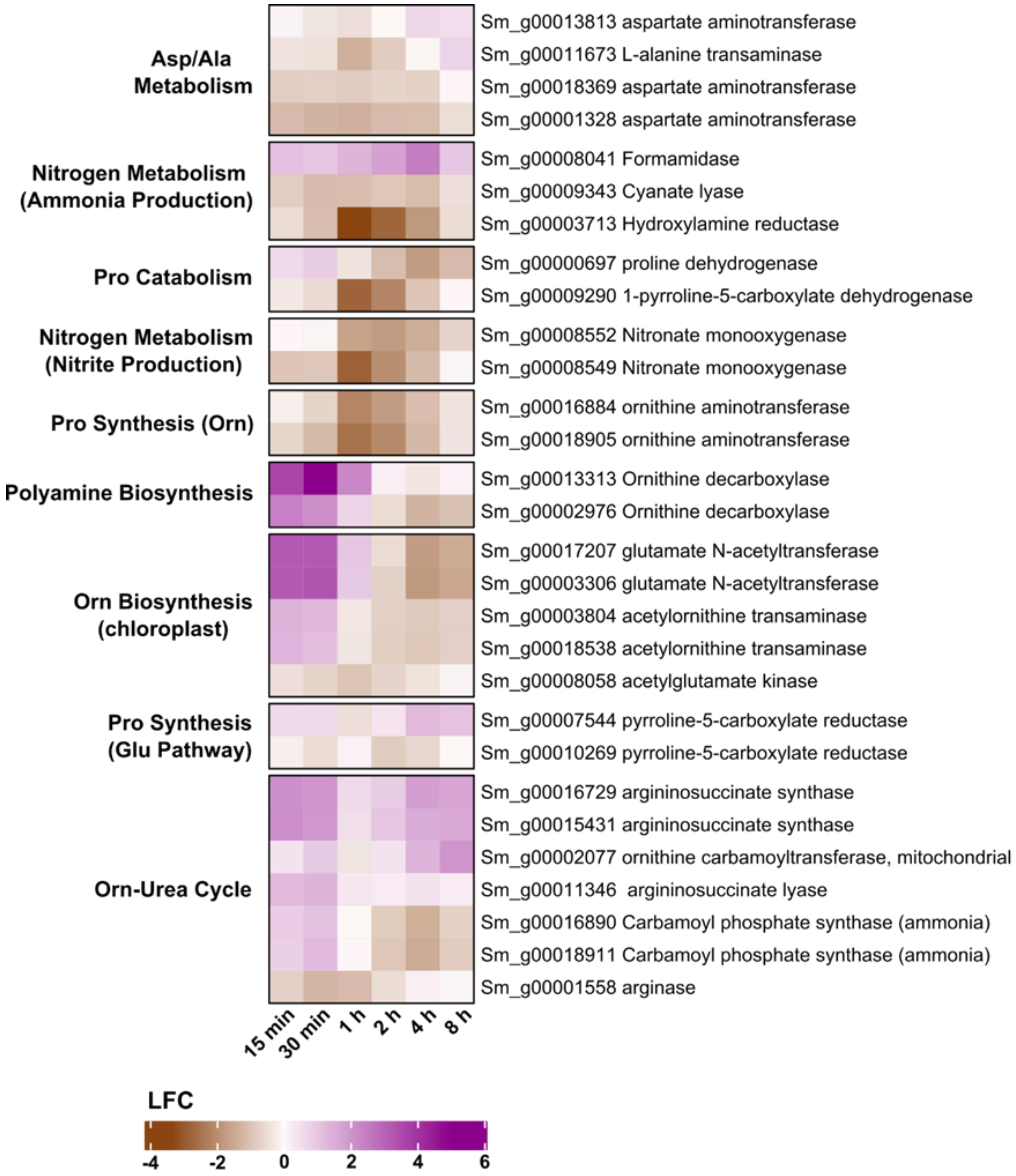


**Supplementary Fig. S3 (continued) | Time-resolved expression of nitrogen metabolism.** Continued from the previous page. Heatmap showing the expression of differentially expressed genes associated with nitrogen assimilation over the 8-hour time series. All shown genes were significant in two or more consecutive time points.

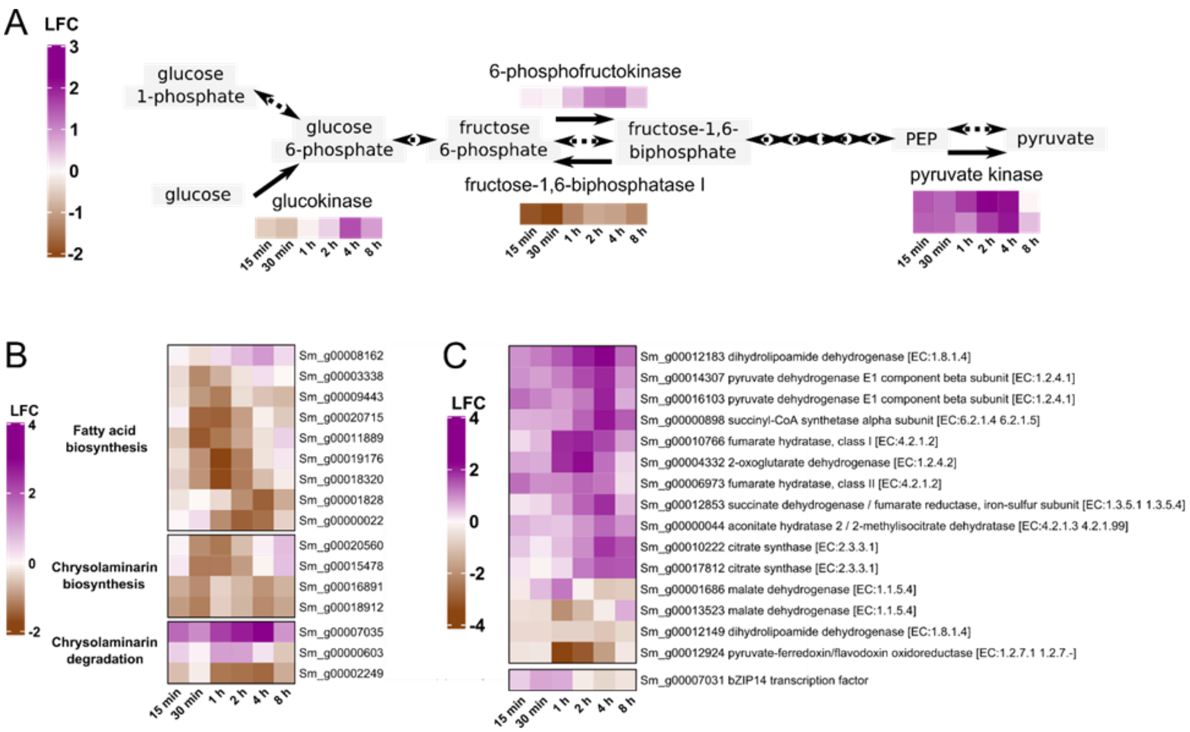


**Supplementary Fig. S4 | Time-resolved expression of carbohydrate and storage molecule metabolism.** Expression of differentially expressed genes involved in carbohydrate metabolism and energy storage over the 8-hour time series. All shown genes were significant in two or more consecutive time points. **a.** Heatmaps of the irreversible steps of glycolysis/gluconeogenesis. Each row is a single gene at each time point. **b.** Heatmap showing expression of differentially expressed genes involved in biosynthesis and degradation of storage molecules. **c.** Heatmap showing expression of differentially expressed genes involved in the TCA cycle, including the *bZIP14* transcription factor which regulates the TCA cycle.

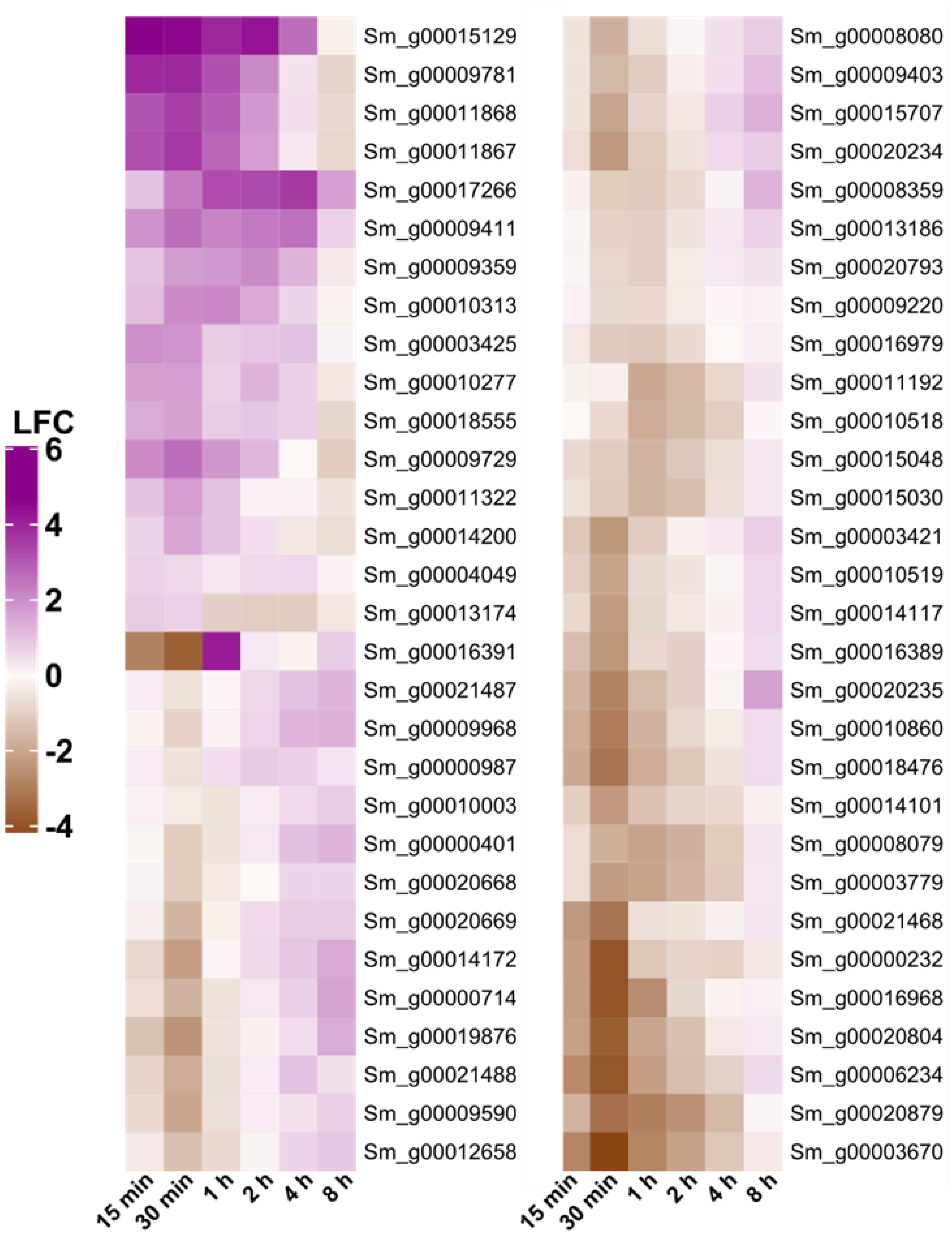


**Supplementary Fig. S5 | Time-resolved expression of genes encoding light harvesting proteins.** Heatmap showing the expression of differentially expressed genes associated with GO:0009765 (photosynthesis, light harvesting) over the 8-hour time series. All shown genes were significant in two or more consecutive time points.

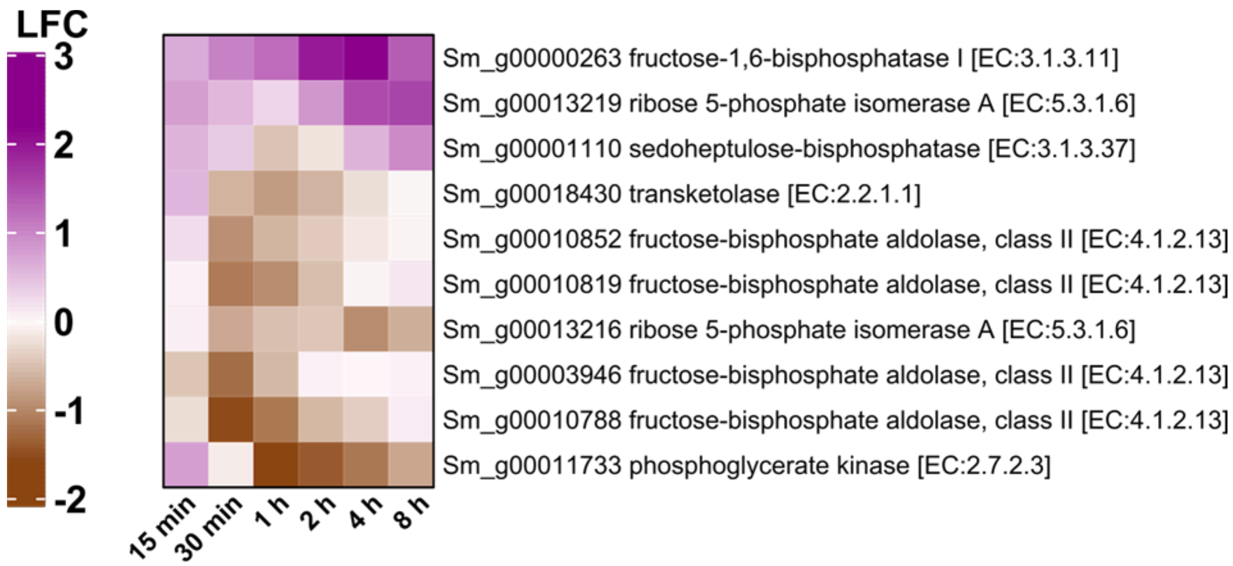


**Supplementary Fig. S6 | Time-resolved expression of genes involved the Calvin cycle.** Heatmap showing the expression of differentially expressed genes from the Calvin cycle over the 8-hour time series. All shown genes were significant in two or more consecutive time points. In case a gene functions in both the cytosol and chloroplast, only homologs localized to the chloroplast were included.

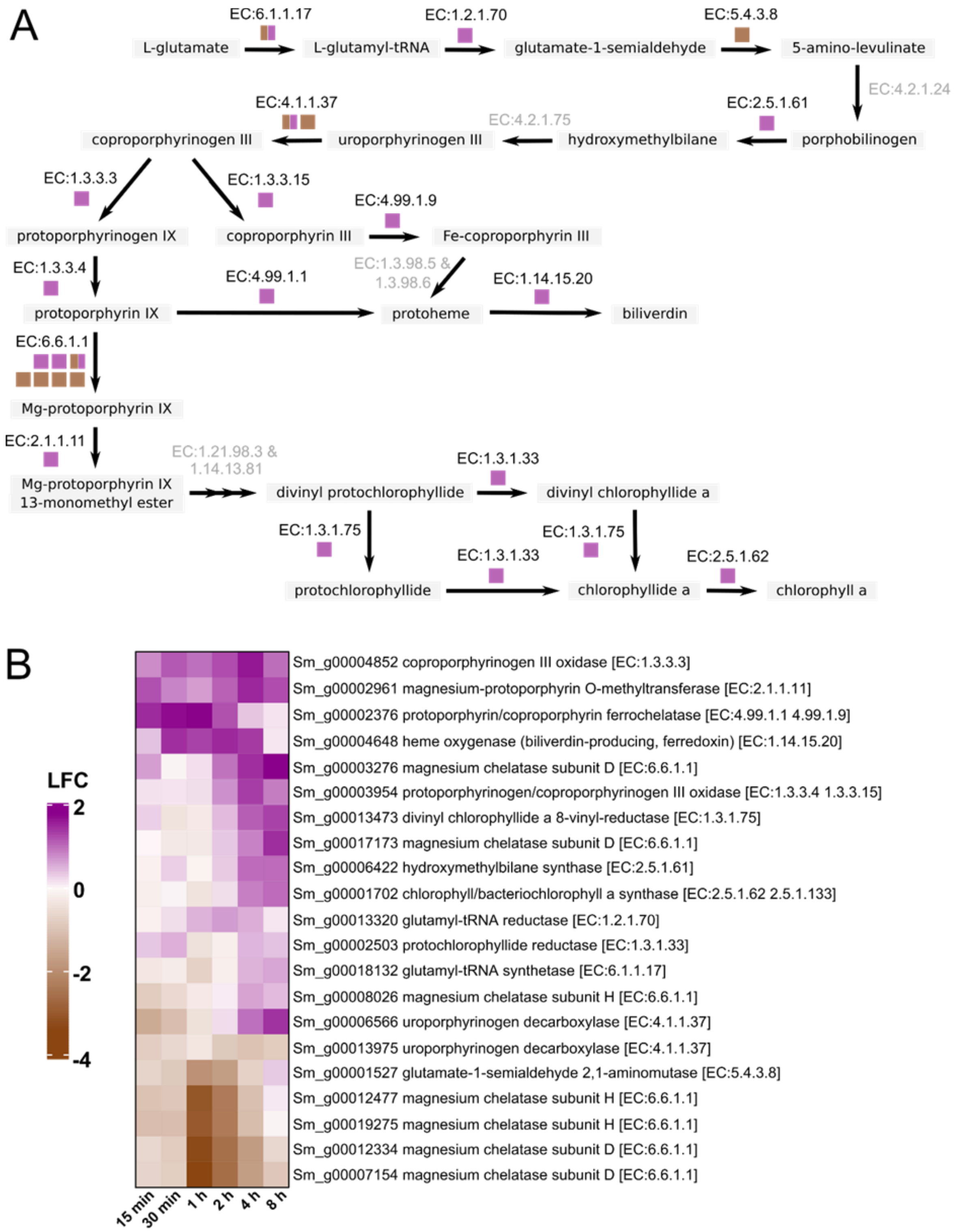


**Supplementary Fig. S7 | Time-resolved expression of genes involved in chlorophyll biosynthesis. a.** Pathway and **b.** Heatmap showing the expression of differentially expressed genes involved in chlorophyll biosynthesis over the 8-hour time series. All shown genes were significant in two or more consecutive time points.

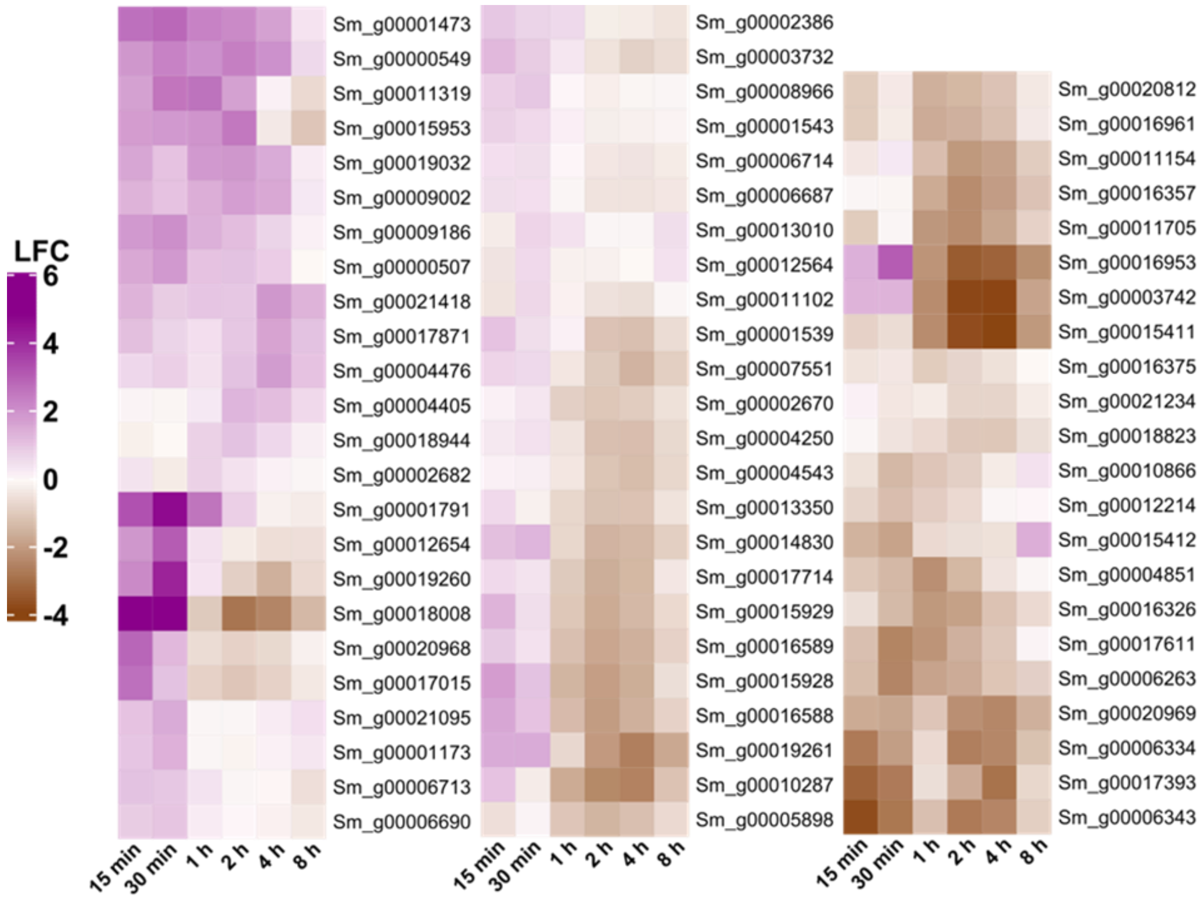


**Supplementary Fig. S8 | Time-resolved expression of heat shock factors/proteins.** Heatmap showing the expression of differentially expressed genes annotated as heat shock factors/proteins over the 8-hour time series. All shown genes were significant in two or more consecutive time points.

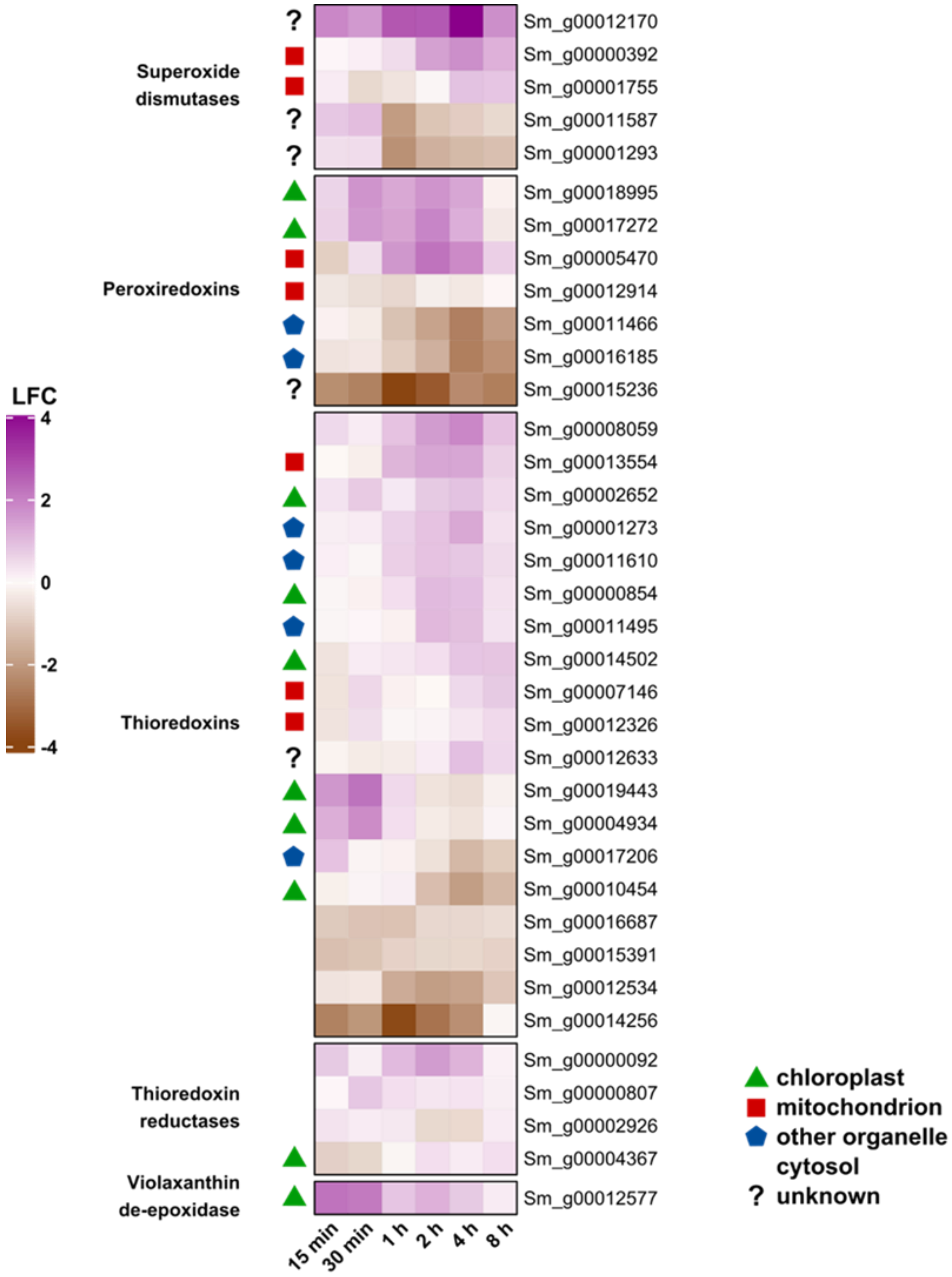


**Supplementary Fig. S9 | Time-resolved expression of genes involved in the mitigation of reactive oxygen species (ROS).** Heatmap showing the expression of ROS mitigation-associated differentially expressed genes over the 8-hour time series. Localization is indicated to the left of each gene. All shown genes were significant in two or more consecutive time points.

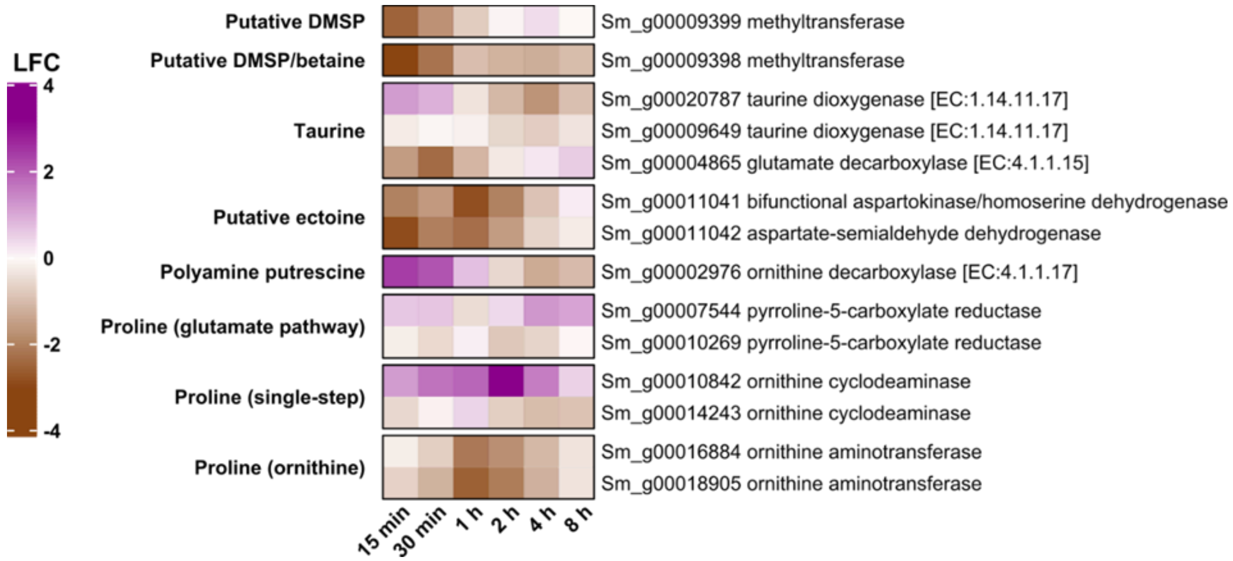


**Supplementary Fig. S10 | Time-resolved expression of genes involved in osmolyte biosynthesis or degradation.** Heatmap showing the expression of differentially expressed genes involved in the biosynthesis and degradation of osmolytes over the 8-hour time series. All shown genes were significant in two or more consecutive time points. The scale indicates logFC.

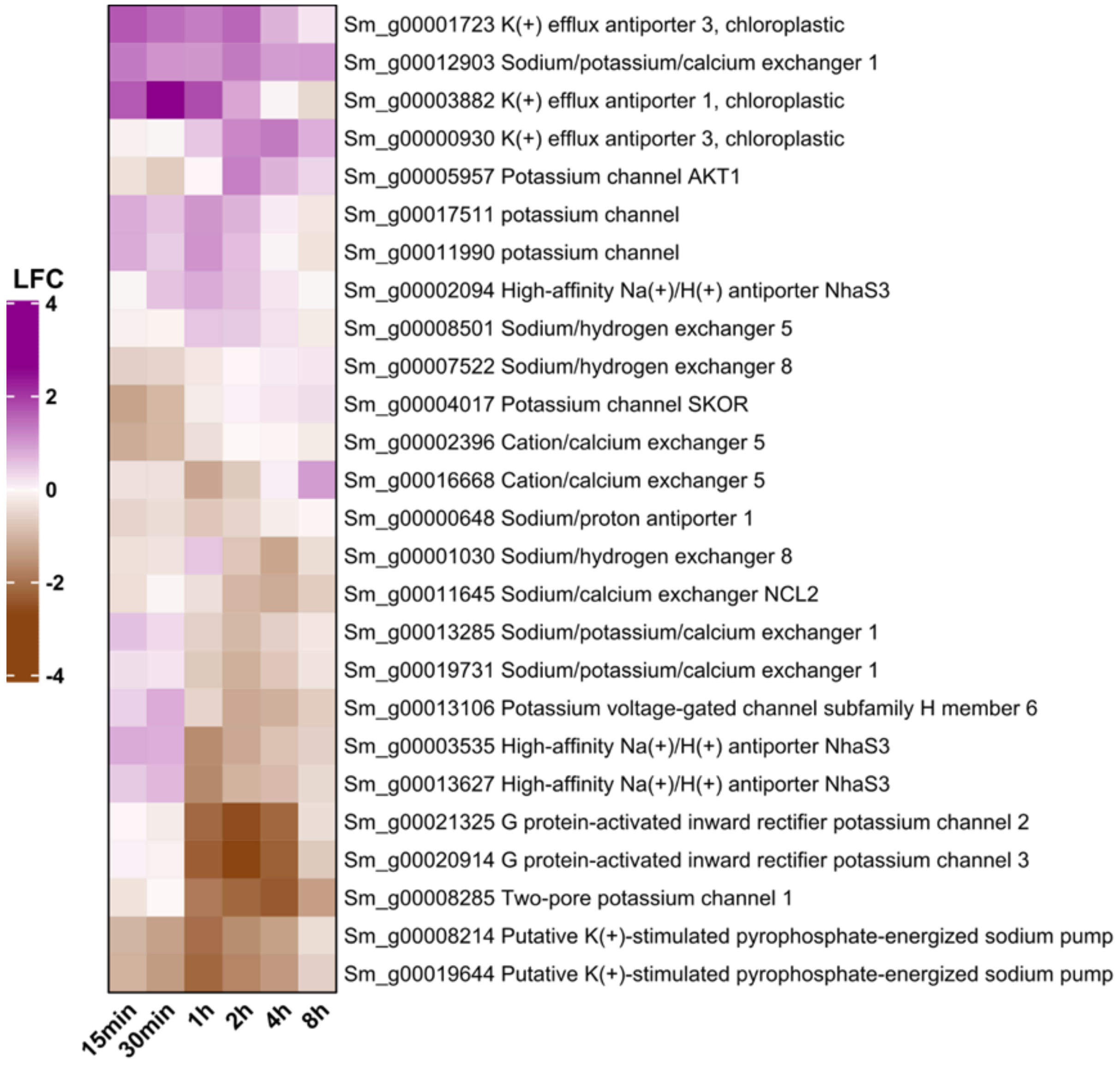


**Supplementary Fig. S11 | Time-resolved expression of ion transport.** Heatmap showing the expression of differentially expressed genes involved in $K^+$ and $Na^+$ ion transport over the 8-hour time series. All shown genes were significant in two or more consecutive time points.

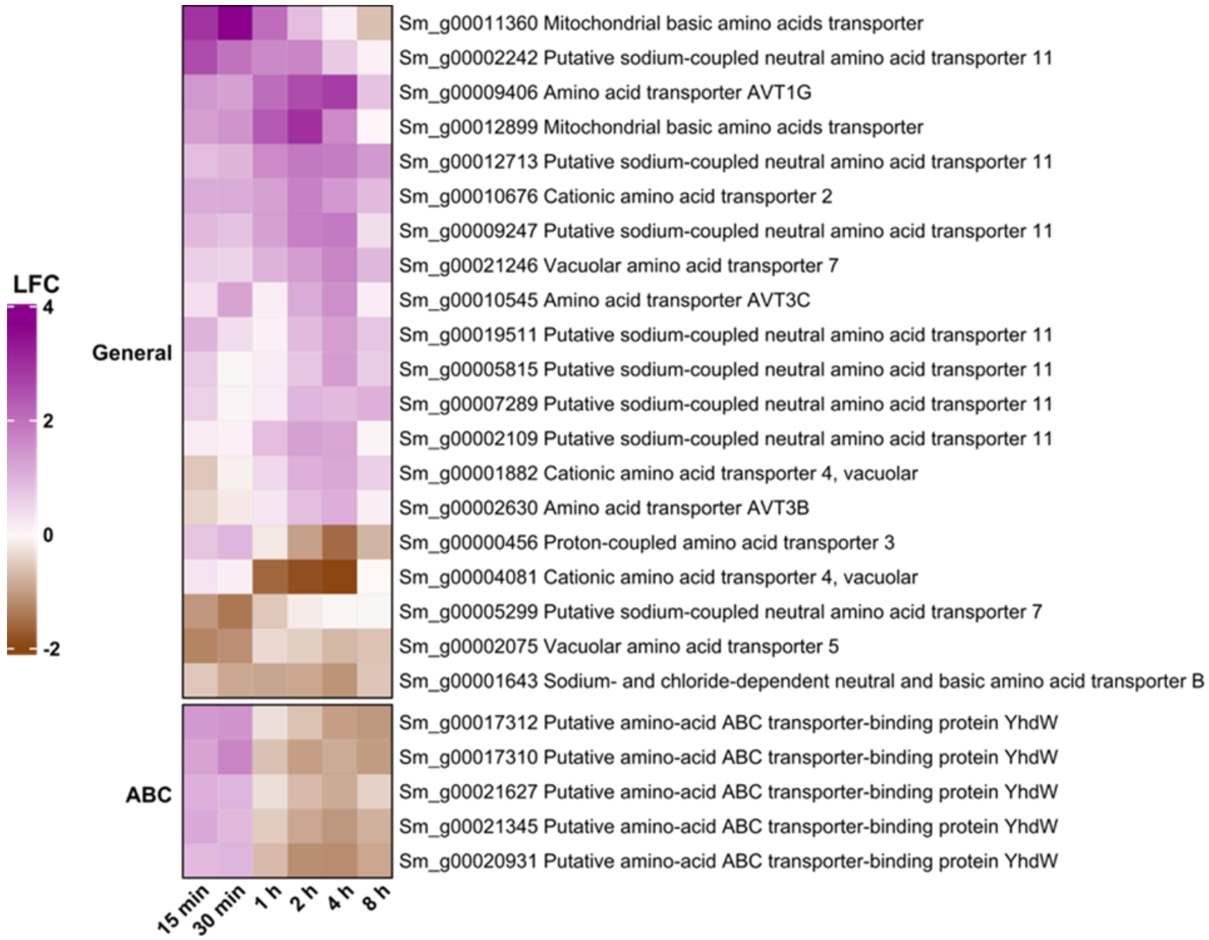


**Supplementary Fig. S12 | Time-resolved expression of amino acid transport.** Heatmap showing the expression of differentially expressed genes involved in amino acid transport over the 8-hour time series. All shown genes were significant in two or more consecutive time points.

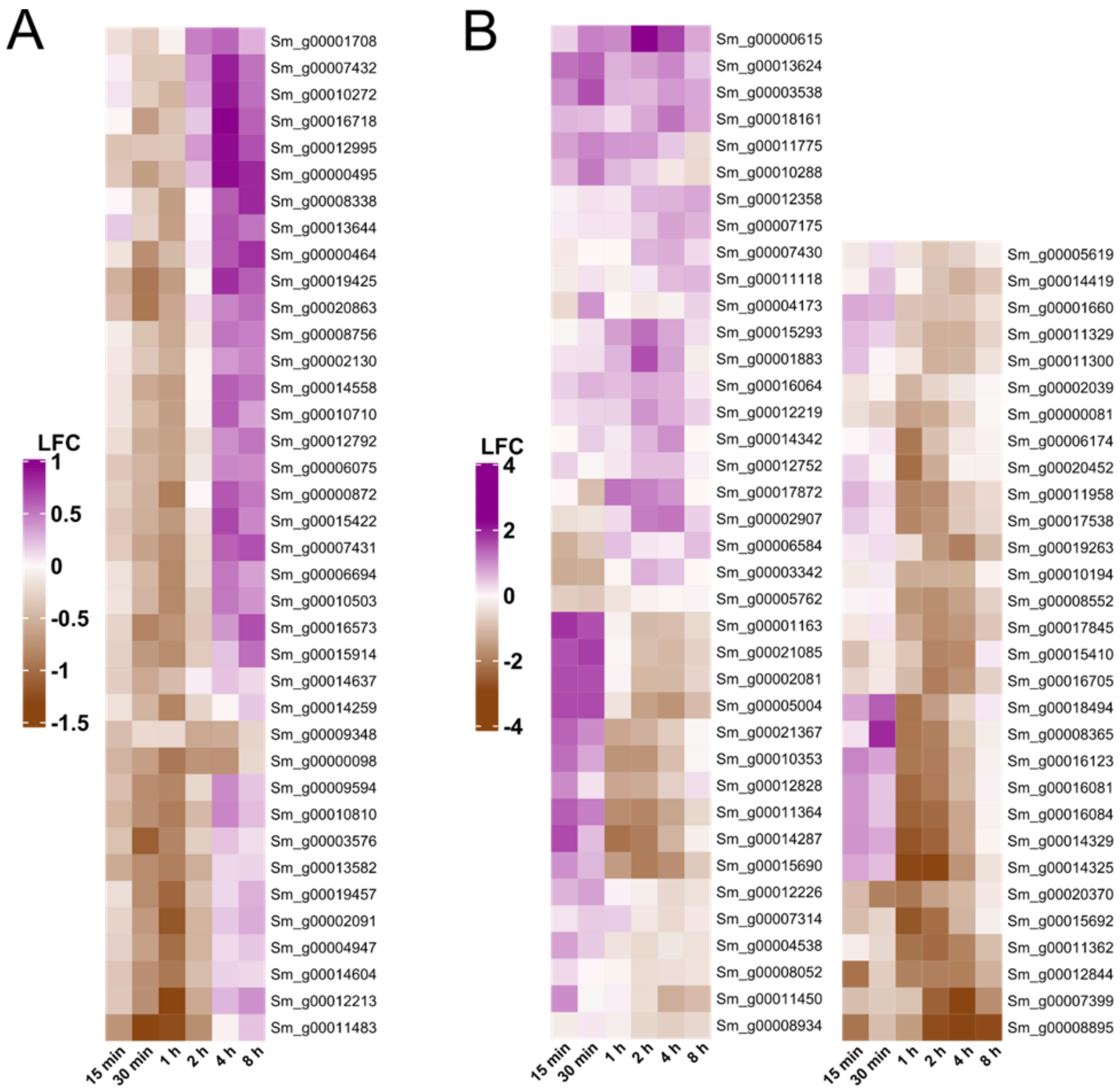


**Supplementary Fig. S13 | Time-resolved expression of proteasome and serine-type endopeptidase activity.** Heatmap showing the expression of differentially expressed genes associated with **a.** proteasome activity and **b.** serine-type endopeptidase activity over the 8-hour time series. All shown genes were significant in two or more consecutive time points.

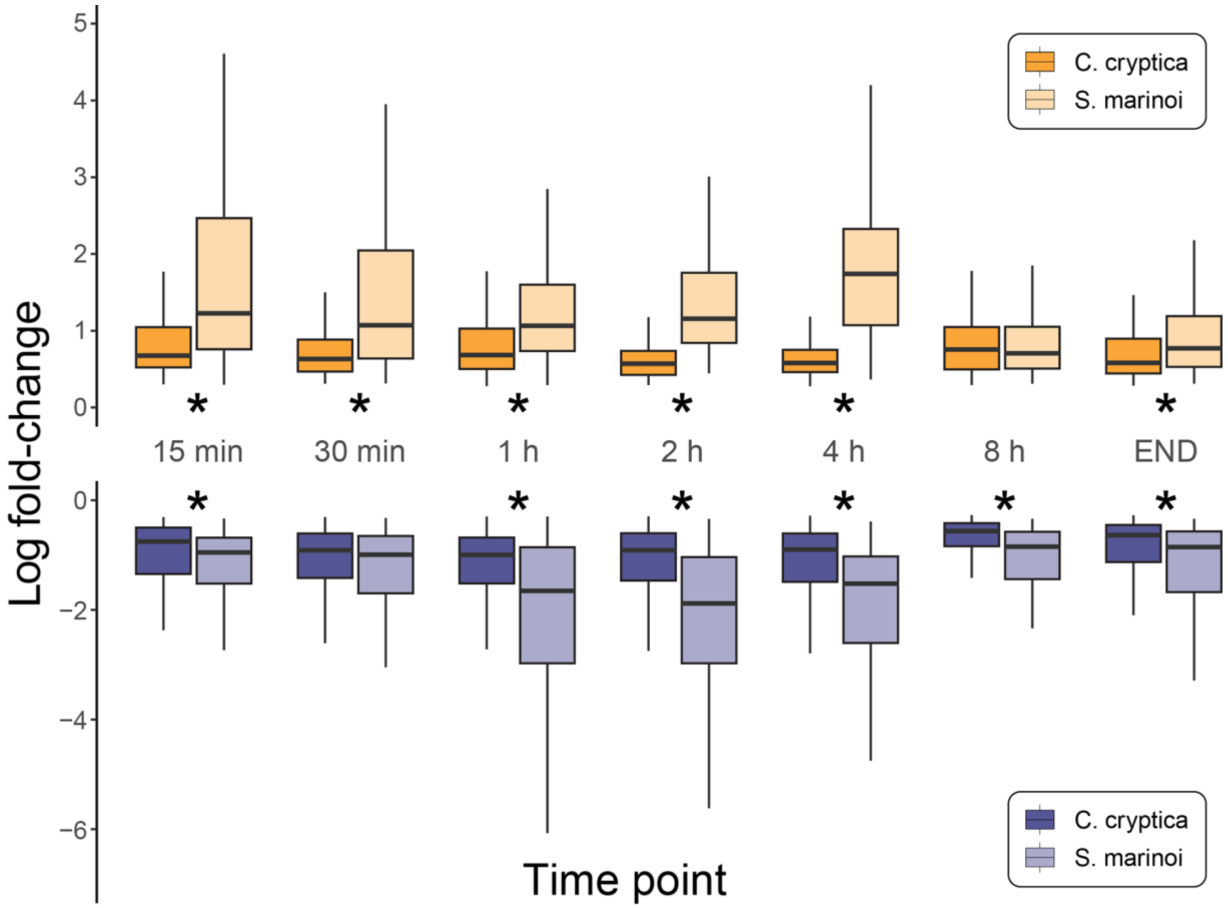


**Supplementary Fig. S14 | Differences in the strength of the hyposalinity stress response.** Distribution of LFC values for all significantly expressed homologs, by direction of expression, in *S. marinoi* (teal) and *C. cryptica* (green). LFC values were summarized per orthogroup to obtain one observation per species. Stars indicate a significant difference (p-value < 0.05) in population mean ranks by a two-sided, two-sample Wilcoxon signed rank test. Outliers not pictured.
